# Spontaneous rhythmic and tool-assisted drumming across variable tempo and technique in a captive chimpanzee

**DOI:** 10.1101/2025.09.26.678729

**Authors:** James Brooks, Vesta Eleuteri, Jelle van der Werff, Shinya Yamamoto

## Abstract

Rhythmic drumming on percussive instruments is a common element of music across human cultures. Chimpanzees also drum on the buttress roots of trees and other resonant objects, and often combine their drumming with long-distance pant-hoot vocalizations. Previous studies have suggested that chimpanzees select drumming substrates for their acoustic properties and that chimpanzee drumming shares core elements of human musicality (e.g., non-random timing, isochronous - metronome-like rhythm, and de-contextualised production in captivity). But to what extent can chimpanzees flexibly control their drumming rhythm across percussive media and techniques? Here, we report on a long-lasting drumming event by a captive chimpanzee named Toon, which was performed across multiple pant-hoot displays and employed diverse action forms, including the use of multiple drumstick-on-drum multi-tools. We find flexible rhythmic production and isochrony across drumming implements, techniques, and tempos. Further, we describe tool set transport and reuse, selectivity of percussive acoustic properties associated with different vocal elements, and variable use of the facial expression “play face” associated with faster and less variable rhythms, potentially indicating intrinsic enjoyment of such rhythms. Together, these findings provide evidence for key elements of human musicality in Toon’s drumming, supporting the hypothesis of shared evolutionary roots of human and chimpanzee drumming.

## Introduction

Music is a cultural phenomenon unique to humans (*Homo sapiens*). To understand the evolutionary origins of music, comparative research therefore focuses on the cognitive and biological capacities that underlie our ability to produce and perceive music, i.e. *musicality* ^1,2^. In the field of biomusicology ^3^, a dominant approach has focused on identifying and studying such capacities in distantly related species (i.e. targeting convergent evolution) ^4–8^, while our nearest relatives, the great apes, have received less attention (i.e. targeting conserved traits) ^9^. Across cultures, humans produce music by drumming on percussive instruments ^10,11^, and while African nonhuman apes also drum on a variety of substrates ^12–15^, the relationship of nonhuman ape drumming to the origins of human musicality remains controversial.

In the wild, chimpanzees (*Pan troglodytes*) drum on resonant tree buttresses generating low-frequency sounds that can travel over a kilometer ^16,17^. Wild chimpanzees mainly drum during traveling and resting contexts ^12,15,18^ but also during agonistic displays ^16,19^. While chimpanzees do not show individual variation when drumming during agonistic displays, they show individual styles (and a potential for signature patterns) when drumming during traveling and resting, potentially to communicate their own location to distant group members and thus facilitate fission-fusion dynamics ^12,15,18^. These findings are suggestive that chimpanzees may be able to flexibly control drumming for different functions. Chimpanzees also show selectivity in their use of percussive surfaces; for example they select larger and thinner buttress roots to drum on ^20,21^, they prefer resonant tree species for both buttress drumming ^20^ and percussive stone throwing ^22^, and they sometimes drum on resonant man-made objects in their environment ^23^. While these findings suggest flexibility in chimpanzee’s drumming behaviour, they do not conclusively demonstrate that individual chimpanzees can control and modulate their drumming.

Chimpanzees often combine drumming with their species-typical “pant-hoot” vocalizations ^15,16^. Pant-hoots are loud calls found across several contexts and are typically composed of four phases ^24^. The “introduction” phase consists of low intensity “hoos”, which can grade into a “build-up” phase characterised by low intensity voiced inhaled and exhaled elements. The build-up usually develops into the “climax” phase, which consists of high-intensity and high-frequency loud variable voiced elements resembling screams. The climax then may grade into the “let-down” phase, in which elements decrease in intensity and frequency, but that is not always included ^15,24,25^. While drumming can occur in each phase, Eleuteri et al. ^12^ reported that most travel and resting drumming was produced during the build-up and climax phases, and showed individual variation in the timing of drumming within pant-hoots. However, the amount of intra-individual flexibility in the combination of pant-hoot and drumming signals has never been assessed empirically.

Rhythm is a core aspect of human music and shows a number of properties that seem universal across cultures ^10,11^. A particularly prominent universal rhythm is isochrony, i.e. the regular, metronome-like spacing of sounds ^10,26^. To date, three studies have demonstrated the presence of isochrony in chimpanzee drumming behaviour ^27–29^. One recent study demonstrated non-random timing and isochrony in wild chimpanzee buttress drumming, as well as subspecies variation in drumming rhythms ^29^. While western chimpanzees (*P. t. verus*) drummed isochronously, eastern chimpanzees (*P. t. schweinfurthii*) drummed by alternating shorter and longer intervals between hits. Relatedly, van der Vleuten et al. ^28^ also recently reported isochrony across both vocal and motoric behaviour of chimpanzee displays in two distinct groups living in captivity, building on a report by Dufour et al. ^27^ (discussed below). Notably, consistent isochrony was found across individuals, with individual chimpanzees showing specific tempo preferences. However, whether individual-specific preferred tempi reflect individual-level constraints (e.g. anatomical, physiological differences) or whether individuals can flexibly modulate form and tempo during isochronous drumming remain to be determined.

To investigate intra-individual variation and flexible control of chimpanzee drumming behaviour, long-lasting events (i.e. a few minutes) are particularly important. Most spontaneous chimpanzee drumming is relatively short-lasting (i.e. a few seconds), thus precluding several important statistical tests at the event level. Only a few notable long-lasting drumming events have been reported. First, Dufour et al. ^27,30^ report a sustained drumming event by a captive adult male chimpanzee in the Netherlands named Barney, who maintained a long bimanual drumming bout of 685 hits for 4 minutes on an upturned barrel. Rhythmic analyses indicated the presence and maintenance of rhythms typical of human percussive music, namely isochrony and binary rhythms (alternating interval durations). Second, Matsusaka et al. ^31^ report on two wild young male chimpanzees in Tanzania, Cadmus and Michio, who drummed repeatedly on clay pots in two separate events. Michio drummed almost 200 hits in 6 minutes using a variety of techniques (i.e. drumming with both hands on different locations and orientations of the pot) and showing a play face, a facial expression used by chimpanzees during play and indicating excitement ^31^. The authors interpreted the event as indicating that Michio was seeking out and enjoying new sounds for their own sake, a crucial feature of human musicality. Finally, Watson et al. ^32^ reported flexible tool-assisted drumming by one captive adult male chimpanzee in the United States named EH. On multiple occasions EH assembled, and even repaired, an innovative multi-material surface to perform drumming during pant-hoots. EH fashioned this surface uniquely to drum, often assembling it even before starting to vocalize, suggesting forward planning of the drumming event.

The previous case reports of a few notable individuals highlight the importance of paying attention to and describing in detail extended events of chimpanzee drumming. However, these events have not yet permitted direct tests of rhythmic flexibility across varied percussive techniques and pant-hoot display phases in chimpanzee drumming. We report here an observation of a captive chimpanzee named Toon engaging in a sustained pant-hoot and drumming acoustic display of over 400 percussive hits in 12 minutes. Toon drummed with his hands, feet, and with a variety of tools, on a variety of surfaces, and across multiple distinct pant-hoots and display phases. We find that this relatively long-lasting drumming event shows flexible rhythmic production (i.e. sustained isochrony across different tempi and motor patterns) and other elements of human musicality reported in prior chimpanzee drumming events (e.g., multi-tool use, presence of play face; ^27,31,32^). We describe and discuss Toon’s overall rhythmicity (i.e. rhythm types and drumming speed), the influence of different pant-hoot display phases, drumming implements and surfaces on drumming rhythm, and the influence of drumming rhythm on play faces.

## Results and discussion

We defined a drumming event as the entire video containing several drumming bouts, an acoustic bout as periods of vocalization and/or drumming separated by periods of rest without any sound production, a drumming bout as a period of drumming with the same implement (limbs or tools) separated by less than 5s, and a drumming hit as an audible sound created by hitting drumming implement against a drumming surface. We followed previous definitions of pant-hoot display phases, and additionally coded limb laterality, drumming implement and surface, and the presence of a play face. See Table S1 for complete description of coding scheme. The beginning of the recording, including the first drumming hits, can be seen in Video 1.

### Drumming production across implements, surfaces, and display phases

Toon used both hands, both feet, two sticks, and one cardboard box as percussive implements (i.e. item used to hit a surface) against nine distinct percussive surfaces (i.e. item hit by the percussive implement). The majority (69.0%) of hits were produced in bouts using hands or feet as the percussive implement. Toon used a tool as a percussive implement in bouts representing 29.4% of hits. The remainder of hits were produced in bouts alternating between limb and tool. See Video 2 for an example of the varied action forms and sound production methods incorporated into Toon’s drumming and pant hoot displays.

Toon’s drumming occurred mostly during the climax (54.5% of hits) and introduction (17.2%) phases of his pant-hoot display, while infrequently during the build-up phase (1.8%). Toon performed only one pant-hoot display with a let-down phase, in which he drummed for five hits (1.2%). 25.3% of his drumming occurred without any associated pant-hoot displays and Toon sometimes drummed before producing any vocalization. Our results differ from the typical pattern for combinations of pant-hoots and drumming in wild chimpanzees reported in Eleuteri et al. ^12^. In that study, wild chimpanzees produced drumming bouts most frequently during the build-up and climax pant-hoot phases, and drumming preceding vocalizations was unreported. However, most of the pant-hoot drums were recorded in the contexts of traveling and resting (See Table 5 in Eleuteri et al ^12^), with only a few observations during agonistic displays.

Toon’s choice of implement and surface for different pant-hoot phases was relatively consistent across the drumming event. Specifically, almost all tool-assisted drumming (both as implement and surface) that was integrated into a pant-hoot display occurred during the introduction phase and was performed from the same elevated platform. During the climax phase, Toon drummed from the ground against the enclosure’s front fence using his arms to produce a loud rattling sound. In the one observed let-down phase, Toon drummed on a surface in a location not otherwise used at any point across the drumming event (i.e. a different area of fence towards the back of his enclosure).

The different acoustic properties of his environment, especially the different use of the louder more resonant sounds of the metal fence in the climax compared to softer sounds produced by tool-assisted drumming in the introduction, suggest that Toon was selectively seeking different percussive sounds for different elements of his acoustic display. Sound selectivity has been shown in the general drumming behaviour of wild chimpanzees ^20–22^, but this report presents the first evidence of a chimpanzee repeatedly, and consistently, incorporating different percussive sounds into different phases of the same acoustic display.

### Drumming rhythmic properties

An inter-hit interval (i.e. IHI) represents the latency between two consecutive drumming hits. Using normalized rhythm ratios of consecutive IHIs ^5^, we compared Toon’s drumming to a random distribution of rhythm ratios where each ratio occurs with uniform probability (Figure 1A; See Supplementary Information for more details). We find that Toon drums with non-random timing (two-sample Kolmogorov-Smirnov *D* = 0.192, *p* < .001). Rather, Toon’s drumming was highly isochronous, meaning that consecutive inter-hit intervals were of equal or similar duration (185 out of 309 ratios were isochronous, 59.9%; Wilcoxon signed-rank test *V* = 136, *p* < .001). Use of isochrony did not differ between introduction and climax phases (Figure 1B; Wilcoxon signed-rank test *W* = 26, *p* = 0.846), nor between tool-assisted and unassisted drumming (Figure 1C; i.e. using only hands and/or feet; Wilcoxon signed-rank test *W* = 32, *p* = 0.503), nor between hands and feet (however the small number of foot drumming hits may have led to loss of statistical power; Figure 1D; Wilcoxon signed-rank test *W* = 39, *p* = 0.101). These results show that Toon creates and maintains isochrony across varied drumming forms and pant-hoot display phases within the same drumming event, demonstrating flexibility in his ability to employ isochronous drumming. This finding shows Toon’s ability to use and maintain regular rhythms regardless of the motor movement employed (i.e. drumming by hands, drumming by feet, or drumming with specific tools).

**Figure 1.**
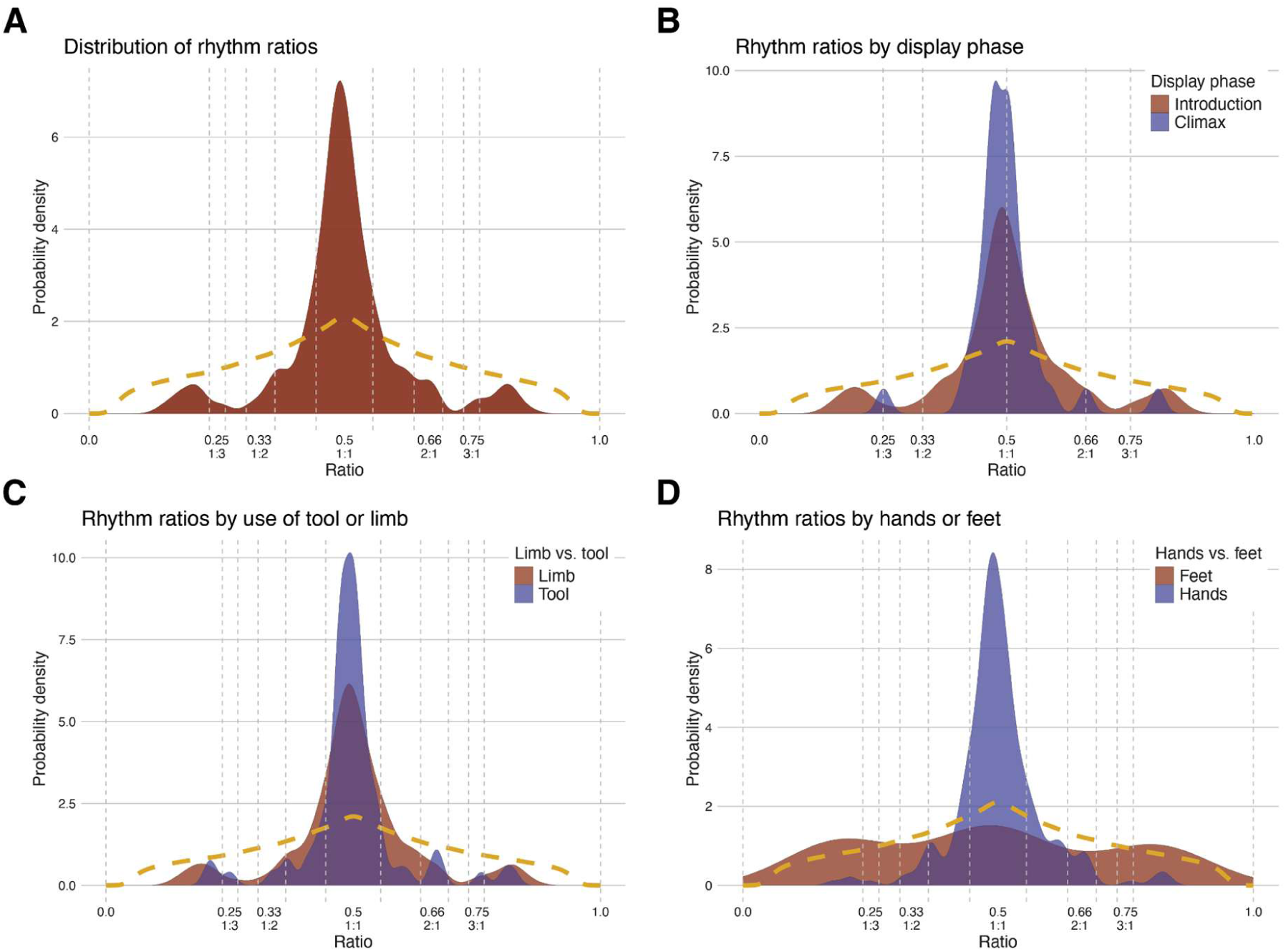
Density distributions of the rhythm ratios of the inter-hit intervals within drumming bouts showing Toon maintains isochrony across drumming implements and display phases. Each rhythm ratio represents the normalized ratio (r) between two consecutive inter-hit intervals (IHI) within drumming bouts, normalized such that their value lies between 0 and 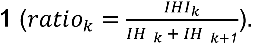 Here, 0.5 indicates a 1:1 relationship (two consecutive intervals of equal duration), 0.33 indicates a 1:2 relationship (an interval followed by one twice its duration), 0.25 a 1:3 relationship (an interval followed by one thrice its duration). ^5^. The dotted yellow line indicates a random distribution of rhythm ratios resulting from uniformly random inter-hit intervals and thus representing what random drumming would look like. **(A) Distribution of rhythm ratios across all drumming bouts.** Toon produced more isochronous inter-hit intervals as can be observed by chance (i.e. in the random rhythm ratio distribution). **(B) Distributions of rhythm ratios by pant-hoot display phase.** Toon showed consistent isochrony regardless of pant-hoot phase. **(C) Distributions of rhythm ratios by use of a tool (e.g. a stick) or limb (feet or hands).** Toon showed consistent isochrony when using tools or limbs. **(D) Distributions of rhythm ratios by use of hands or feet.** Toon did not significantly differ in rates of isochrony when using hands compared to feet, however few drumming events produced with his feet limit statistical power.

In addition to isochrony, in one drumming bout Toon seemed to create and sustain a complex rhythm where he repeatedly used a short-long-long pattern of consecutive IHIs by holding the fence’s ceiling with his arms and swinging left to right, hitting the side fence with two hits (one for each foot) on his left and just one hit on his right (Figure 2; Video 3). This rhythm is consistent with the binary rhythms reported by Dufour et al. ^27^ for Barney, achieved there through bimanual drumming, and suggests that Barney is not entirely unique in his rhythmic abilities. Towards the end of this swinging bout, Toon held the fence’s ceiling with only one hand and continued swinging back and forth, now tapping the fence with the other hand in between alternating foot hits. This three-limb rhythmic pattern was not sustained long enough to examine any repeated patterns. Nonetheless, the two limb pattern indicates the ability to create sustained repeating patterns with varied hit spacing within a largely isochronous drumming event.

**Figure 2.**
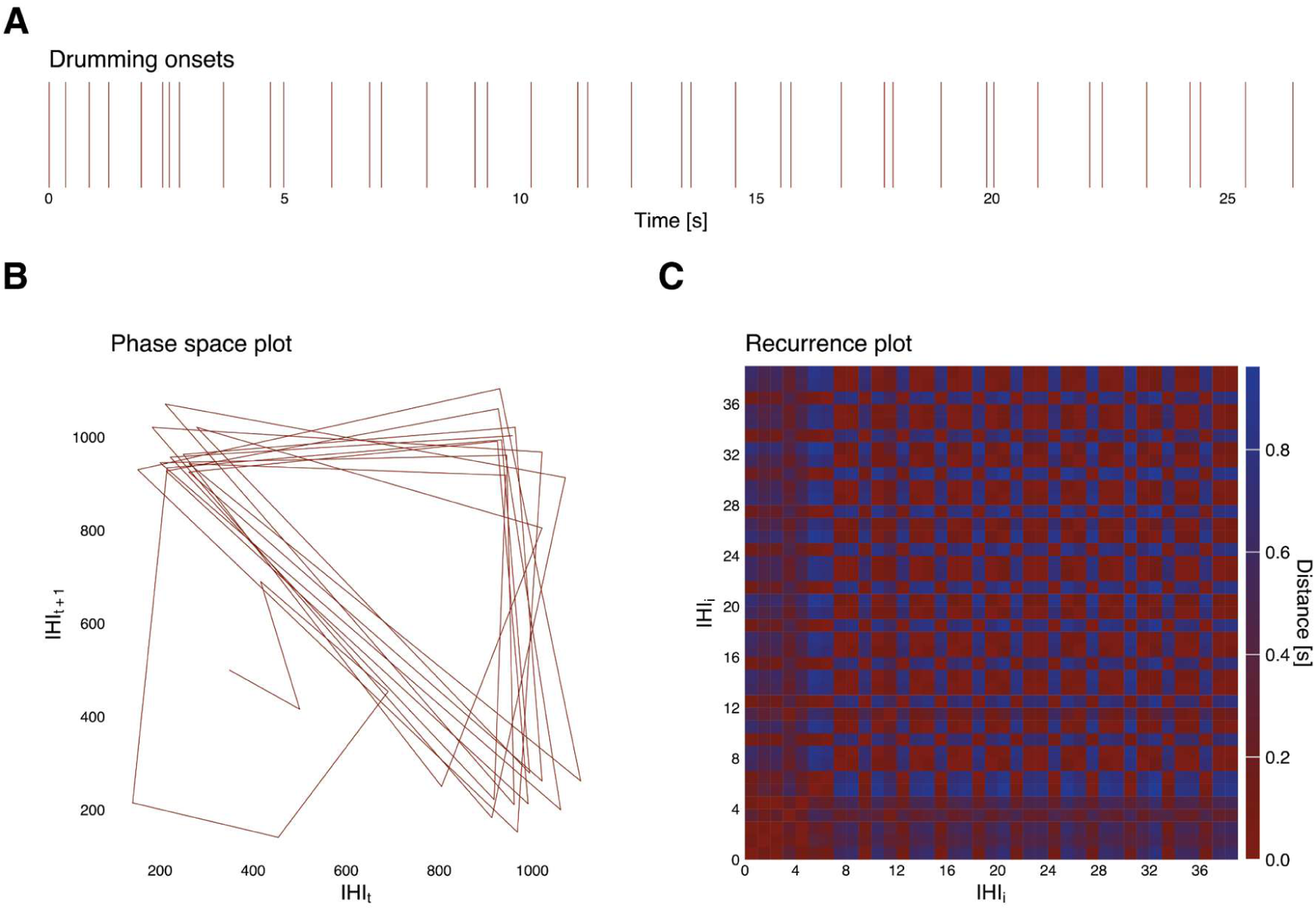
Rhythmic visualizations of the fourth drumming bout showing a short-long-long pattern. **(A) Event plot illustrating the fourth drumming bout**. Each vertical line represents one drumming hit onset. **(B) Phase space plot showing the fourth drumming bout inter-hit intervals (IHIs)**. The phase space plot has the duration of the IHI on the *x* axis and the duration of its consecutive IHI on the *y* axis ^38^. Each point in the plot thus represents a pair of consecutive IHIs. Lines between these points are drawn in order of appearance in the drumming bout. The geometrical pattern here shows a drumming pattern consisting in the repeated use of short–long–long inter-hit intervals. **(C) Recurrence plot showing a repeated pattern within the fourth drumming bout.** Recurrence plots illustrate relative similarity and difference between the durations of all pairs of IHIs (i.e. the distance between them), which here provides a visualization of repetition in rhythmic patterns. The colour indicates how similar (red) or different (blue) the duration of pairs of IHIs are from each other, while the position of each IHI is indicated on both axes. The pattern that emerges (i.e. a grid pattern) here indicates the repeated use of a short–long–long IHI drumming pattern.

Isochrony is a central property of human musicality ^33,34^ that allows us to synchronize behaviour during collective music and dance, and has been thus linked to social cohesion and coordination in humans ^26^. This drumming event indicates that our capacity to dynamically sustain isochrony across production forms ^26^ is shared with chimpanzees and raises questions about the evolutionary emergence of these capacities. It should also be noted that Toon is a western chimpanzee, and that a recent study found isochronous drumming in western chimpanzees but not in their eastern counterparts ^29^, highlighting that evolutionary considerations are in crucial need of greater reporting on other subspecies. Future studies should investigate the contextual and social drivers of isochronous drumming by chimpanzees and pay attention to potential subspecies differences to further our understanding of the origins of music.

### Drumming tempo across acoustic phases, drumming implements, and rhythms

Overall, the mean IHI was 501.27 ms (i.e. around 100 bpm), which is slightly faster than typically reported in human spontaneous tapping experiments (∼600ms [22]) but slower than reported in wild chimpanzee drumming (∼229ms ^29^), though similar to that reported in a direct comparison of tapping in both humans and chimpanzees (400 to 600 ms for both species) ^35^. Drumming during the introduction phase was slower (*M* = 825 ms) than during the climax phase (*M* = 431 ms; *t*(231) = 10.20, *p* < .001, 95% CI of β [1.17, 1.73]). While both phases showed strong isochrony, the average drumming speed during the climax phase was roughly double that of the introduction phase (Figure 3A), suggesting that faster drumming may be due to increased arousal during the climax phase compared to the introduction phase of the display. Toon drummed slower with tools (*M* = 696 ms) than with his limbs (*M* = 431 ms; *t*(311) = 9.093, *p* < .001, 95% CI of β [0.81, 1.26]), and slower with his feet (*M* = 664 ms) than with his hands (*M* = 479 ms; *t*(311) = -4.241, *p* < .001, 95% CI of β [-1.06, -0.39]). Drumming with limbs was faster than drumming with tools, likely due to more frequent hand-drumming in the climax phase (see below). However, it is worth noting that there are two clear clusters both with and without tools (Figure 3B), suggesting that Toon is capable of modulating his drumming speed using limbs and tools. Considering his strong use of isochronous drumming across drumming implements mentioned above, these results show that Toon is able to maintain isochrony at varied tempi across percussive techniques within a single extended drumming event. Consistent isochrony alongside variable drumming speed indicates substantial rhythmic flexibility not previously reported in chimpanzees.

**Figure 3.**
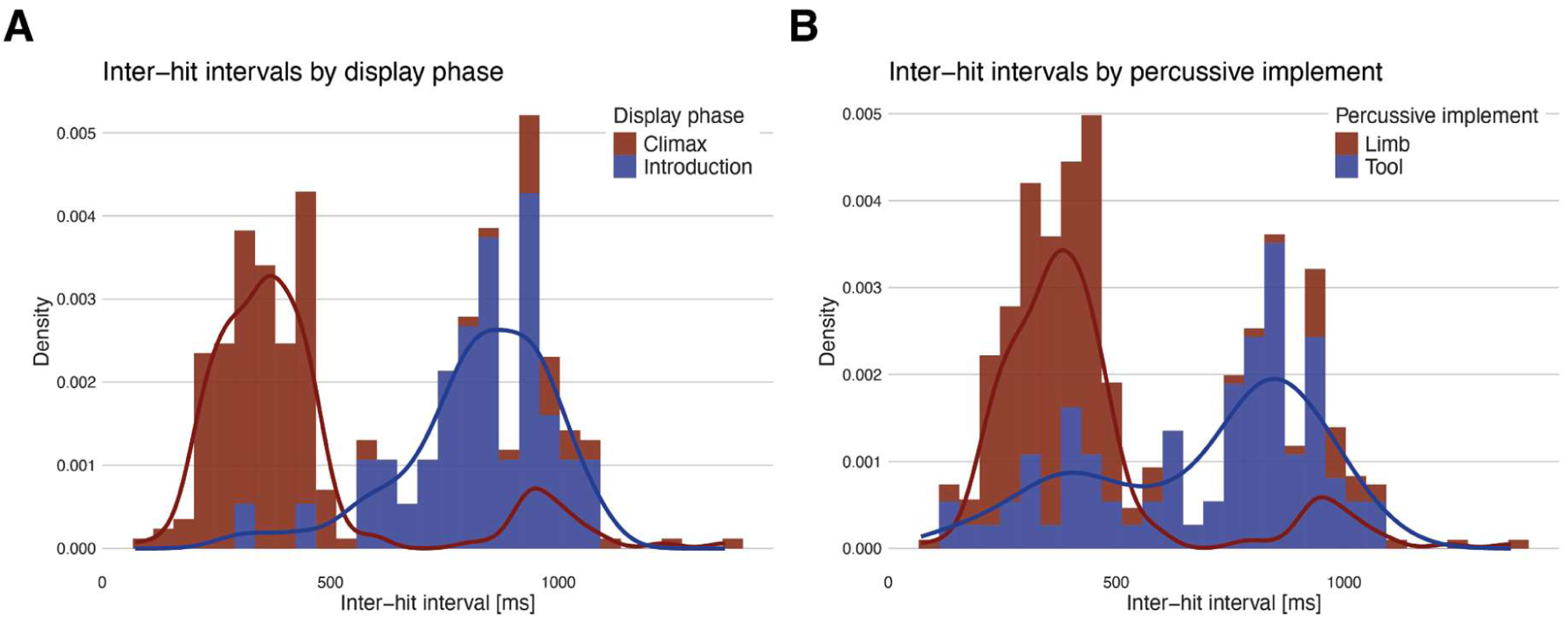
Histograms and density curves of inter-hit intervals (IHIs) across all drumming bouts. **(A) IHIs by phase of the pant-hoot display**. Drumming during the introduction phase was around twice slower than drumming during the climax phase. **(B) IHIs by use of limb (feet or hands) or tool (e.g. a stick).** Drumming performed with tools was around twice slower than drumming performed with the limbs.

### Use of play face during drumming

We tested whether play faces were associated with particular rhythmic properties of Toon’s drumming bouts (Figures 4 & 5; Video 4). Specifically, we compared the mean IHI and variability of IHIs accompanied by a play face and IHIs without a play face (see Supplementary Information). Drumming with play face was faster (*M* = 374 ms) than drumming without play face (*M* = 598 ms; *t*(311) = -8.534, *p* < .001, 95% CI of β [-1.08, -0.68]). Variability of IHIs was also lower when the play face was present (*SD* = 91 ms) than when it was not present (*SD* = 295 ms; *F*(177) = 10.55, *p* < .001, 95% CI [7.64, 14.45]). Thus, play faces were associated with faster and less variable drumming inter-hit intervals. This may be related to Toon’s increased arousal with faster and more predictable rhythms. In other words, Toon’s play faces may be suggestive of his enjoyment of fast and predictable drumming, where he can best “feel the groove”. This is consistent with Matsusaka’s ^31^ report of a young wild chimpanzee playing by creating sound, and may underlie Toon’s intrinsic motivation to drum for self-amusement, a core property of human music.

**Figure 4.**
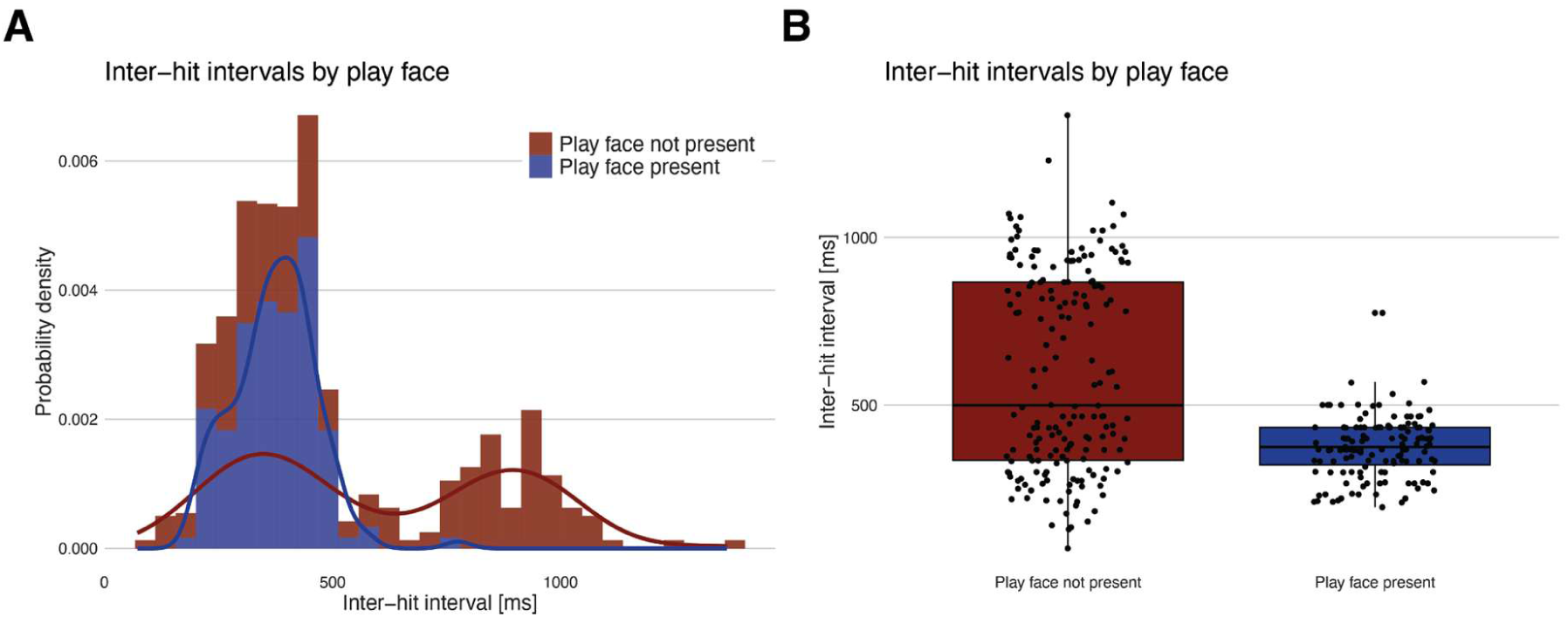
Plots of drumming inter-hit intervals (IHIs) according to use of play face. **(A) Histograms and density curves of IHIs according to whether the play face was present or not present**. Drumming was faster when Toon drummed with a play face as compared to without. **(B) Boxplots showing the dispersion of IHIs according to whether the play face was present or not present.** Each dot represents one IHI. There was less variability in the distribution of IHIs when the play face was present compared to when it was not present.

**Figure 5.**
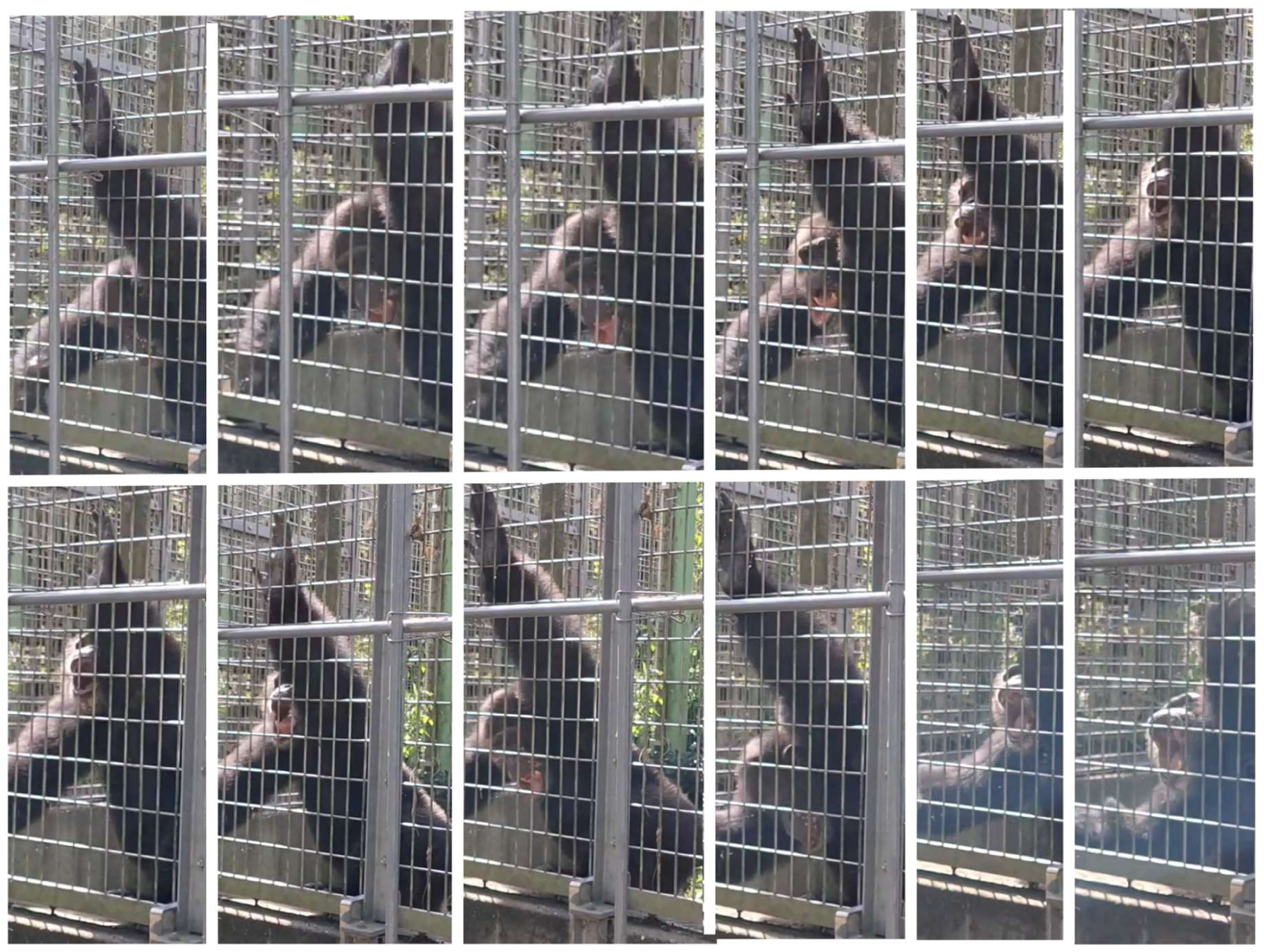
Example photos showing the “play face” facial expression that was associated with faster drumming (shorter inter-hit intervals).

### Tool and multi-tool drumming techniques

Toon used at least two distinct techniques of tool use for sound production. These include the “drumstick” technique, where he held one of the two sticks and hit the percussive surface, and a “dropping” technique, where he repeatedly picked up and dropped the cardboard box tool to create sound. Some of the tools he used as percussive surfaces (i.e. one stick and the box) were also used as percussive implements, indicating flexible use of each tool as both surface and implement for sound production. The variety of objects used and of techniques used for the same objects, for no clear purpose besides the production of percussive sounds, indicates sound selectivity and flexibility in Toon’s tool-assisted drumming display.

Toon also showed use of tool composites in bouts where he used one tool (the “drumstick”) to hit another tool (the “drum”) (see Video 2; Figure 6). Additionally, in some bouts, Toon sequentially hit multiple different percussive surfaces with the same tool similarly to humans drumming on a drumkit. Multi-tool use has been previously reported in a captive chimpanzee drumming event by Watson et al. ^32^. However, the multi-tool “instrument” reported there is a tool composite of an entirely different kind: the chimpanzee in Watson et al. ^32^ used a soft material to cushion a harder plastic tub which he then hit with his hands. Here, Toon used a stick to hit a percussive surface directly. A previous study showed use of sticks to perform drumming in palm cockatoos ^6^. To our knowledge, this is the first report of a non-human animal spontaneously using tools as both drums and drumsticks as well as the first primate to use a “drumstick” drumming technique.

**Figure 6.**
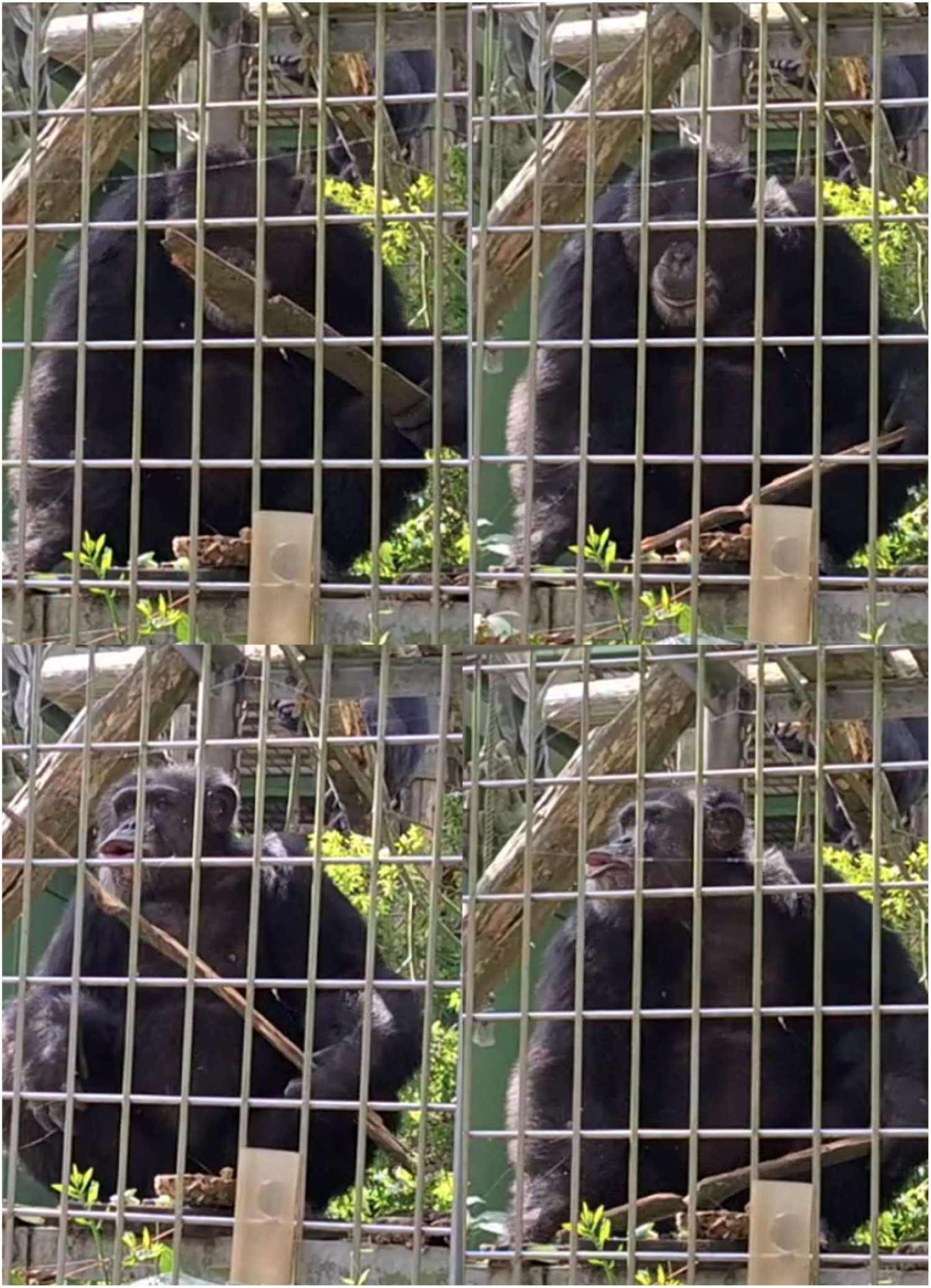
Example photos showing one of the drumstick-on-drum tool sets used by Toon.

One important factor that we could not explicitly account for here is the origin of Toon’s “drumstick” drumming technique (i.e. the use of a drumstick tool for drumming). While we have no reason to suspect any specific trained behaviour or exposure to similar behaviour from humans, it is impossible to rule out human influence. However, observations of Toon’s groupmates suggests that drumming with “drumstick” tools may be socially transmissible. This drumming technique has been observed in 6 out of 13 individuals in Toon’s group, but in none of the other chimpanzees or bonobos groups at Kumamoto Sanctuary (>50 apes) and has not been described in other reports. The individuals in Toon’s group have different backgrounds and life histories. For example, one of the 6 individuals observed drumming with a drumstick, Shoubou, lived in isolation undergoing biomedical experiments for most of his life before arriving at Kumamoto Sanctuary. Thus, the most plausible explanation for the use of “drumstick” drumming exclusively in this group is social influence. However, for nearly all the other individuals, “drumstick” drumming has been observed once or twice. For Toon, tool-assisted drumming (including the drumstick technique) is observed on a more regular basis and can happen up to several times per month. However, its use is unpredictable and usually lasts only a few drumming hits, making systematic and rhythmic analyses of other drumming events challenging. Similarly, drumming by hand is observed more frequently across more individuals, especially against the resonant metal fence, but is difficult to predict and often short-lasting, making tests of flexibility in the expression of drumming rhythms across the chimpanzee group challenging.

### Forward planning and flexibility of tool-based drumming

Toon used only three distinct tools as percussive implements (two sticks and one cardboard box) but used each several times, even retrieving and reusing the same tools multiple times for different drumming bouts (e.g. Figure 7, Video 2). Toon simultaneously transported both the drum and drumstick to his preferred drumming location (i.e. the elevated platform), marking a novel case of tool set transport for acoustic signal production. Toon typically retrieved tools after tossing them away during the building excitement of a previous pant-hoot (i.e. during the build-up and climax phases). He consistently carried the same tools back to the same elevated platform, where he drummed during all introduction phases of his display. At one point, Toon also fashions a tool by stripping another stick of its leaves and twigs, however he did not use this tool for drumming (see Video 2). It should be noted that, within the enclosure, there were several other sticks and objects available, including some closer to him than those retrieved, but Toon consistently sought the same tools for drumming. The retrieval and reuse of the same tools may indicate some level of forward planning and, thus, intentionality ^27,32^ in his drumming, especially given his tendency to bring the tools to a preferred drumming location.

**Figure 7.**
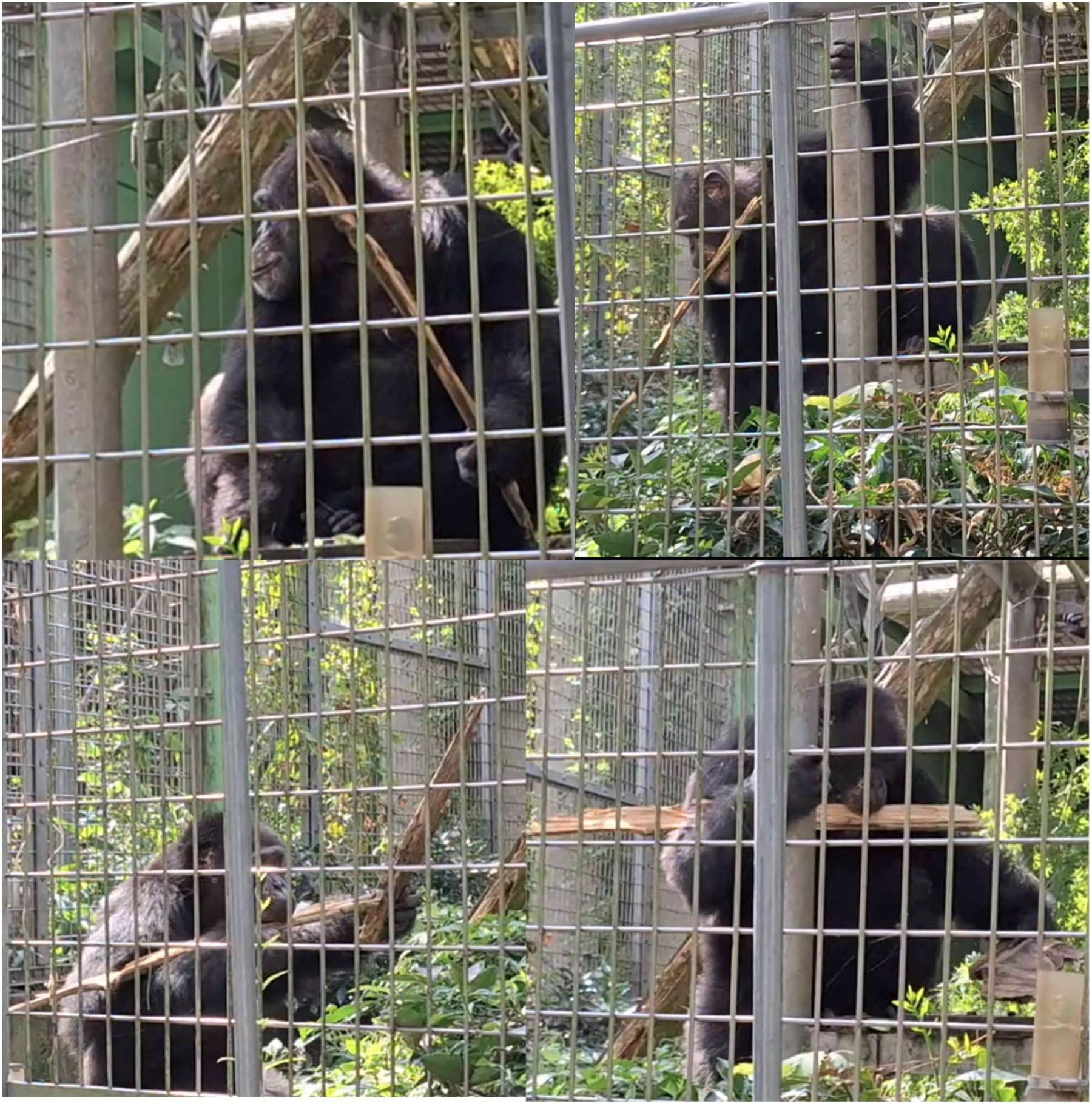
Example photos showing tool set transport.

### Drumming limb laterality

Toon seemed to show considerable left-biased laterality in his drumming, both while using tools and while using limbs (84.8% of drumming hits were produced using a left-side limb, while 12.9% were produced using a right-side limb, with the remaining 2.3% produced bilaterally). There were insufficient right-limb hits to assess potential differences in rhythmicity and isochrony associated with laterality. Across populations, evidence suggests a right-handed bias in chimpanzees, although it is less pronounced than in humans ^36,37^. Notably, Michio, the young chimpanzee in Matsusaka ^31^ who showed extended drumming with play faces, also predominantly drummed with his left handHandedness was not reported in Watson et al. or Dufour et al. ^27,32^ Remaining agnostic on any conclusions from this limited case, we encourage researchers exploring the origins of music to pay attention to chimpanzees’ laterality in drumming production.

## Conclusion

We presented here a detailed analysis and description of a particularly noteworthy event of extended tool-assisted rhythmic drumming by one captive male chimpanzee. We found strong use of isochronous rhythm across different percussive techniques, tempo variability while maintaining stable rhythm, drumming sound selectivity by context (pant-hoot display phase), as well as flexible use and transport of tools and multi-tools. We also show presence of play face associated with fast, less variable rhythms, possibly indicating enjoyment of quicker and more predictable rhythms ^31^, evidence of forward planning in tool transport suggesting intentional production of drumming ^27^, and use of multiple drumstick-on-drums tool sets integrated into the same acoustic display as a drum “kit”. Our study extends upon the findings by Dufour et al. ^27^ on Barney’s spontaneous rhythmic drumming and those by Watson et al. ^32^ showing multi-tool drumming in EH. We suggest it may be Toon’s unique motivation for drumming, rather than any particularly unique ability, which permitted us to detect these findings showing several key elements of human musicality. We encourage other researchers across sites to report on “exceptional drummers” so that future works might be able to identify what factors may lead to these rare but geographically widespread cases of ape drumming with music-like characteristics. A deeper understanding of great ape drumming behaviour will be crucial to future comparative research on the evolution and conserved roots of human musicality, and our findings suggest the ability to create and maintain regular rhythm across varied tempo and motor movement, including through the use of tools, can be found in our closest relatives.

## Author contributions

Conceptualisation: JB, VE, JW; Data collection: JB; Methodology: JB, VE; Formal analysis: JW; Supervision: SY; Writing - Original Draft: JB, VE, JW; Reviewing and editing: JB, VE, JW, SY.

## Data availability

All data and scripts for their analysis are available from GitHub.

## Acknowledgements

We thank Toon, the other chimpanzees, and all of the care staff at Kumamoto Sanctuary for making this study possible. We thank the Yamamoto and Ravignani labs for valuable discussions. We also thank Kumamoto Sanctuary/Etsuko Nogami for sharing the photograph of Toon used in the graphical abstract. JB was funded by the Japan Society for the Promotion of Science (24KF0058), VE was funded by the Austrian Science Fund (FWF) “DK Cognition and Communication 2”: W1262-B29 [10.55776/W1262] awarded to the Department of Behavioral and Cognitive Biology of the University of Vienna, JW was funded by the European Union (ERC, TOHR, 101041885, awarded to Andrea Ravignani), and SY was funded by Japan Society for the Promotion of Science (KAKENHI 24H02200) and JST FOREST Program (JPMJFR221I).

## Supplemental videos

All supplemental video files can be found at: https://drive.google.com/drive/folders/1Ebljg-Ia2QOEjdgA5GBc3JiUeIhOBONr?usp=sharing

## Captions

**Video 1. Video showing the beginning of the recording including the first drumming hits and vocalizations of the drumming event.**

**Video 2. Video of Toon’s pant-hoot display showing the several elements reported here, including drumstick-on-drum tool set transport and use, integration of drumming across pant-hoot phases, variable use of percussive media by pant hoot phase, and variable expression of “play face”.**

**Video 3. Video of the fourth drumming bout showing a short-long-long pattern.**

**Video 4. Video of Toon showing the “play face” facial expression that was associated with faster drumming (shorter inter-hit intervals).**

