## Supplementary Information for "Spontaneous rhythmic and tool-assisted drumming across variable tempo and technique in a captive chimpanzee"

### Extended methods

#### *Study site and subject*

Kumamoto Sanctuary is a sanctuary for chimpanzees and bonobos in southern Japan. Toon is a Western chimpanzee (*Pan troglodytes verus*) who was born in captivity at Amersfoort Zoo, the Netherlands. He was human-reared for one year before moving to Noichi Zoo, Japan, where he lived until his arrival at Kumamoto Sanctuary in 2008. He has no history in entertainment. Life-history details can be found at shigen.nig.ac.jp/gain/ with GAIN ID 0411. During discussions with care staff about the possible origin of his interesting drumming behaviour it was suggested that it emerged only upon arrival to Kumamoto, possibly due to the fences being particularly resonant.

#### *Data collection*

This observation took place in October 2023 at Building 1 of Kumamoto Sanctuary, where 13 adult male chimpanzees (including Toon) live in a simulated fission-fusion environment where subgroup compositions change on a daily basis. On the day of this recording there was a “birthday party” for a chimpanzee in the neighbouring subgroup (with groupmates familiar to Toon), entailing a larger than average amount of shareable fruit and several familiar human observers outside the enclosure. During the party, Toon was socially housed in a typical subgroup with other familiar males. Since the neighbouring enclosures had the gate closed on this day, Toon could observe but not join the birthday party.

Video recording began when the observer noticed Toon had picked up two sticks and the observer suspected he may begin drumming with them. No other drumming by Toon was observed in the preceding 15-20 minutes, though some brief hand-drumming by others was observed during the birthday party.

*Video coding*

The video was coded with the video coding software Elan 6.2 [1]. To quantitatively analyze Toon's drumming event, we used a detailed coding scheme with each behaviour of interest. Table 1 provides definitions for each coded behaviour. We measured the onset of each hit by combining video playback and visual inspection of the spectrogram (Figure S1).

| | |
|---|---|
| Acoustic bout | From the first vocalization or percussive hit until the last, where bouts are separated by periods of rest without any vocalizations or percussive hits. |
| Display phase | Time subdivision of acoustic bouts into introduction, buildup, climax, letdown. These phase divisions follow pant hoots (see Arcadi [2], Fedurek et al., [3]) and were generalized here into the overall acoustic bout (i.e., including drums in the absence of vocals) for acoustic bouts that themselves contained a pant hoot. Specifically, drum hits prior to the pant hoot build-up were labelled "introduction", during the pant-hoot build-up labelled "build-up" and drums following the peak of a build-up hoot were labelled "climax". A "let-down" phase was only coded when there were decelerating drums following a "climax". In acoustic bouts lacking a pant-hoot, no phase was assigned. |
| Drumming bout | Bout of drumming separated by less than 5s or using a different drumming implement. |
| Drumming event | The entire observed drumming event. |
| Drumming hit | Audible sound created by hitting a drumming implement (e.g., tool, hand) against a drumming surface. Measured at the time of contact. |

| | |
|---|---|
| Drumming implement | The object that moved to make a percussive hit (whether part of the body or an external tool). “Hands” and “Feet” are considered distinct implements. If there is alternation between multiple implements, this alternation is identified as a distinct drumming implement (e.g., stomping a foot between stick drum hits has the implement “stick+feet”). The implement is specific to a drumming bout. |
| Drumming surface | Where the drumming implement makes contact in producing a drumming hit. The surface can vary within a drumming bout (e.g., when playing a drumkit). |
| IHI | Inter-hit interval measures the amount of time between the onset of two successive drumming hits. |
| Laterality | Whether the implement was/was controlled by a limb on the left or right (from Toon’s perspective). |
| Play face | Whether or not a play face (open mouth and loose lips) was displayed (coded for each drumming hit). |

Table S1: Definition scheme used for video analysis.

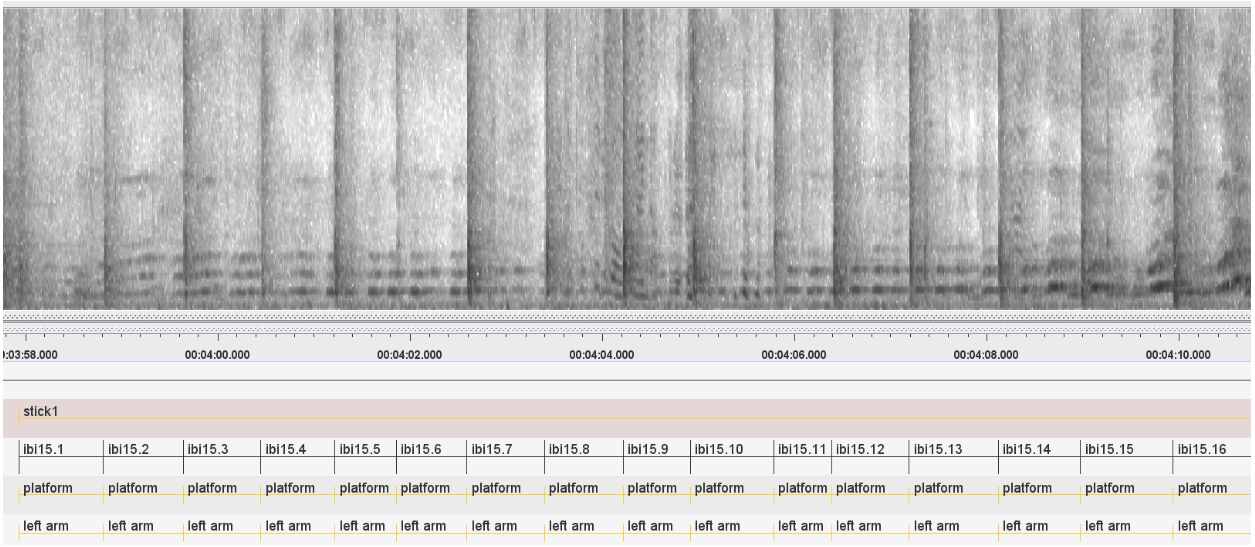


Figure S1: Illustration of spectrogram and coding.

*Supplemental videos:*

*All supplemental video files can be found at:*

*https://drive.google.com/drive/folders/1Ebljg-Ia2QOEjdgA5GBc3JiUeIhOBONr?usp=sharing*

*Captions:*

*Video 1. Video showing the beginning of the recording including the first drumming hits and vocalizations of the drumming event.*

*Video 2. Video of Toon's pant-hoot display showing the several elements reported here, including drumstick-on-drum tool set transport and use, integration of drumming across pant-hoot phases, variable use of percussive media by pant hoot phase, and variable expression of "play face".*

*Video 3. Video of the fourth drumming bout showing a short-long-long pattern.*

*Video 4. Video of Toon showing the "play face" facial expression that was associated with faster drumming (shorter inter-hit intervals).*

*Statistical analysis*

All statistical analyses were based on either inter-hit intervals (IHIs) or on rhythm ratios between adjacent IHIs. IHIs were calculated as the intervals between consecutive drumming hit onsets. Rhythm ratios were then calculated by dividing each inter-hit interval by the sum of itself and the following interval using the formula $ratio_k = \frac{IHI_k}{IHI_k + IHI_{k+1}}$ (Roeske et al., [4]). For the rhythmic analyses reported below we excluded all factor levels (e.g. “fence” from factor “percussive surface”) with fewer than 10 IHIs or 9 ratios. IHIs were used to compare drumming tempo and variability, while rhythm ratios were used to investigate the use of non-random timing and isochrony (following Eleuteri et al.[5]). To investigate tempo differences between different drumming behaviours (e.g., techniques, media, implements, phases), we used one-way ANOVAs with IHI as the dependent variable. To compare drumming variability across different drumming behaviours, we used F-tests with IHI as the dependent variable.

To investigate non-random timing in Toon’s drumming, we compared the distribution of Toon’s rhythm ratios to a simulated distribution of rhythm ratios derived from a uniformly random distribution of 100,000 IHIs (i.e., random distribution). The two distributions of rhythm ratios were then compared using a two-sample Kolmogorov-Smirnov test. The random uniform distribution of IHIs had the shortest observed IHI in Toon’s drumming as the lower bound of the distribution, and the longest observed IHI as the upper bound of the distribution (following Eleuteri et al. [5]).

To test whether Toon drummed more isochronously than can be expected by chance, and to investigate whether Toon drummed (more) isochronously during certain behaviours, we largely followed the established method to test for isochrony from Roeske et al. [4]. However, we incorporated recent suggestions made in Jadoul et al. [6] and recently implemented by Eleuteri et al. [5], using the following method: First, we calculated the normalized rhythm ratios for each pair of IHIs within each drumming bout using the aforementioned formula ($ratio_k = \frac{IHI_k}{IHI_k + IHI_{k+1}}$). Note that these ratios are normalized in the sense that they have a minimum value of 0, and a maximum value of 1. Each of the resulting values represents the durational relationship between two consecutive IHIs; for example, if the second IHI in a pair is twice as long as the first IHI we have a relationship of 1:2, represented as the normalized ratio 0.33, a relationship of 1:1 is represented as 0.5, etc. We then binned the distribution of ratios according to the bin boundaries as defined in Roeske et al. [4]. Then, to specifically test whether Toon drummed more isochronously than chance level, we counted how many of Toon's rhythm ratios fell into the isochronous bin (i.e., 1:1) and compared Toon's isochrony probability to the chance level probability that rhythm ratios derived from the random distribution fell into the isochronous bin (i.e., chance level isochrony probability). Toon's isochrony probability was calculated for each bout as $\frac{number\ of\ isochronous\ ratios\ in\ bout}{total\ number\ of\ ratios\ in\ bout}$. Chance level isochrony probability was calculated using the formula provided in Jadoul et al. (2025), which results in a chance level for the isochronous bin of 0.218. In other words, 21.8% percent of rhythm ratios can be expected to be isochronous when assuming random drumming. We used one-sample Wilcoxon signed-rank tests to compare Toon's isochronous drumming with chance level isochrony.

All statistical analyses were performed in *R* [7]. Analysis scripts and results are provided (see Data Availability). These documents also contain more detailed descriptions of the analyses, as well as examples of how the procedure worked.

1. Wittenburg P., H. Brugman, A. Russel, *et al.* 2006. ELAN : a professional framework for multimodality research. In 1556–1559.
2. Arcadi A.C. 1996. Phrase structure of wild chimpanzee pant hoots: Patterns of production and interpopulation variability. *American Journal of Primatology* **39**: 159–178. https://doi.org/10.1002/(SICI)1098-2345(1996)39:3<159::AID-AJP2>3.0.CO;2-Y
3. Fedurek P., A.M. Schel & K.E. Slocombe. 2013. The acoustic structure of chimpanzee pant-hooting facilitates chorusing. *Behavioral Ecology and Sociobiology* **67**: 1781–1789. https://doi.org/10.1007/s00265-013-1585-7
4. Roeske T.C., O. Tchernichovski, D. Poeppel, *et al.* 2020. Categorical Rhythms Are Shared between Songbirds and Humans. *Current Biology* **30**: 3544-3555.e6. https://doi.org/10.1016/j.cub.2020.06.072
5. Eleuteri V., J. van der Werff, W. Wilhelm, *et al.* 2025. Chimpanzee drumming shows rhythmicity and subspecies variation. *Current Biology* **35**: 2448-2456.e4. https://doi.org/10.1016/j.cub.2025.04.019
6. Jadoul Y., T. Tufarelli, C. Coissac, *et al.* 2025. Hidden assumptions of integer ratio analyses in bioacoustics and music. . https://doi.org/10.48550/arXiv.2502.04464
7. R Core Team. 2019. R: A Language and Environment for Statistical Computing. *R Foundation for Statistical Computing*.
