## Supplementary material for "Spontaneous rhythmic and tool-assisted drumming across variable tempo and technique in a captive chimpanzee": stats-Toon.html

Toon drumming


### Toon drumming

###### Jelle van der Werff

#### 2025-09-17

- Introduction
- Preliminaries
- Data
  cleaning
- Ratios
  statistics method (following Jadoul et al., 2024)
- Descriptive statistics
- Does Toon drum
  rhythmically?
- Does drumming medium
  change the rhythm?
- Does the use of
  hands or feet change the rhythm?
- Does Toon show limb
  laterality?
- Does the used surface
  change the rhythm?
- Does the display phase
  change the rhythm?
- Does the
  presence of a play face change the rhythm?
- Description of drumming bout
  4

### Introduction

Here we present the analysis and results for the drumming data
collected from Toon. We refer the reader to the accompanying publication
XYZ.

### Preliminaries

#### Load packages and data

```
## Loading required package: pacman
```

```
## here() starts at /Users/jellevanderwerff/james_drumming_chimp
```

### Data cleaning

- We exclude the factor levels with fewer than 10 IOIs or 9
  ratios.
- We exclude all factor levels in percussive\_surface that contain a
  ‘/’ (i.e. not the same surface used for all onsets).

```
# For factors percussive_medium, tool_or_limb, percussive_surface, display_phase and play_face, exclude factor levels with fewer than 10 IOIs or 9 ratios
iois.clean <- plyr::ddply(
    iois, c("percussive_medium", "tool_or_limb", "percussive_surface", "limb", "display_phase", "play_face"),
    function(x) {
        if (nrow(x) >= 10) {
            return(x)
        }
    }
)

# For factors percussive_medium, tool_or_limb, percussive_surface, display_phase and play_face, exclude factor levels with fewer than 9 ratios
ratios.clean <- plyr::ddply(
    ratios, c("percussive_medium", "tool_or_limb", "percussive_surface", "display_phase", "play_face"),
    function(x) {
        if (nrow(x) >= 9) {
            return(x)
        }
    }
)

# Exclude all factor levels in percussive_surface that contain a '/'
iois.clean <- iois.clean[!grepl("/", iois.clean$percussive_surface), ]
ratios.clean <- ratios.clean[!grepl("/", ratios.clean$percussive_surface), ]

write.csv(iois.clean, here("data", "processed", "drumming_iois_clean.csv"), row.names = FALSE)
write.csv(ratios.clean, here("data", "processed", "drumming_ratios_clean.csv"), row.names = FALSE)
```

### Ratios statistics method (following Jadoul et al., 2024)

```
bins <- list(
    "1" = c(0, 1 / 4.25), # 1 / 4.5
    "2" = c(1 / 4.25, 1 / 3.75), # 1 / 4 > 1:3
    "3" = c(1 / 3.75, 1 / 3.25), # 1 / 3.5
    "4" = c(1 / 3.25, 1 / 2.75), # 1 / 3 > 1:2
    "5" = c(1 / 2.75, 1 / 2.25), # 1 / 2.5
    "6" = c(1 / 2.25, 1 - 1 / 2.25), # 1 / 2 > 1:1
    "7" = c(1 - 1 / 2.25, 1 - 1 / 2.75), # 1 - 1 / 2.5
    "8" = c(1 - 1 / 2.75, 1 - 1 / 3.25), # 1 - 1 / 3 > 2:1
    "9" = c(1 - 1 / 3.25, 1 - 1 / 3.75), # 1 - 1 / 3.5
    "10" = c(1 - 1 / 3.75, 1 - 1 / 4.25), # 1 - 1 / 4 > 3:1
    "11" = c(1 - 1 / 4.25, 1) # 1 - 1 / 4.5
)

breaks <- c(0, 1 / 4.25, 1 / 3.75, 1 / 3.25, 1 / 2.75, 1 / 2.25, 1 - 1 / 2.25, 1 - 1 / 2.75, 1 - 1 / 3.25, 1 - 1 / 3.75, 1 - 1 / 4.25, 1)

labels <- names(bins)
```

#### Introduction

Different from how integer ratios have previously been normalized, we
here adhere to the new recommendations from Jadoul et al. (2014).

According to Jadoul et al. (2024), the assumption of a uniformly
random distribution of IOIs as the null-hypothesis means that the ratios
are distributed according to the distribution plotted below. The exact
shape of this distribution is dependent on the left and right bounds of
the random uniform distribution used for sampling the IOIs. However, the
shape will remain similar to the one plotted below.

As left (a) and right (b) bounds of the uniform distribution of IOIs,
in our analysis we use the minimum observed IOI duration and the maximum
observed IOI duration, respectively.

The dotted lines in the plot are the histogram bins, following Roeske
et al. (2020), which represent the integer ratios 1:3 (bin centered
around 0.25), 1:2 (bin centered around 0.33), 1:1 (isochrony; bin
centered around 0.5), 2:1 (bin centered around 0.66), and 3:1 (bin
centered around 0.75). The bins in between these integer ratios are the
off-integer bins.

```
integer.ratios.labels <- as.data.frame(list(
    x = c(0.25, 0.335, 0.5, 0.665, 0.75),
    y = c(0.1, 0.1, 0.1, 0.1, 0.1),
    label = c("1:3", "1:2", "1:1", "2:1", "3:1"),
    size = c(4, 6, 10, 6, 4)
))

ggplot(ratios.simulated, aes(x = ratio)) +
    geom_density(linewidth = 2) +
    theme_toon() +
    geom_vline(xintercept = breaks, linetype = "dashed", linewidth = 0.5, colour = "grey") +
    labs(x = "Ratio", y = "Probability density", title = "Random ratios distribution", subtitle = "Assumes uniformly random IOIs") +
    geom_text(
        data = integer.ratios.labels, aes(x = x, y = y, label = label), size = integer.ratios.labels$size,
        family = "Helvetica", inherit.aes = FALSE, show.legend = FALSE
    )
```

#### Preparations

##### Coding ratios categories

We follow Roeske et al. (2020) in how to define the bins for the
integer ratios (see above).

```
ratios.clean$bin <- cut(ratios.clean$ratio, breaks = breaks, labels = labels)
ratios.simulated$bin <- cut(ratios.simulated$ratio, breaks = breaks, labels = labels)
```

##### Surface areas of the bins

According to Jadoul et al. (2024), in order for us to later normalize
the observed counts (seeing as the ratios are not uniformly
distributed), we need to calculate the surface areas of the bins. Here,
we estimate them numerically by counting the number of ratios in each
bin in the simulated data, though it is also possible to calculate them
based on the formula provided by Jadoul et al. (2024), which we do
later.

```
ratios.normalization <- data.frame(bin = as.character(1:11))
ratios.normalization$surface_area <- NA
for (i in 1:11) {
    bin <- bins[[as.character(i)]]
    ratios.normalization$surface_area[ratios.normalization$bin == as.character(i)] <- length(ratios.simulated$ratio[ratios.simulated$ratio >= bin[1] &
        ratios.simulated$ratio < bin[2]]) / length(ratios.simulated$ratio)
}

ratios.normalization$bin_left <- sapply(bins, function(x) x[1])
ratios.normalization$bin_right <- sapply(bins, function(x) x[2])
ratios.normalization$bin_size <- ratios.normalization$bin_right - ratios.normalization$bin_left
ratios.normalization$bin_name <- c("1/4.5", "1/4", "1/3.5", "1/3", "1/2.5", "1/2", "1 - 1/2", "1 - 1/3", "1 - 1/3.5", "1 - 1/4", "1 - 1/4.5")
ratios.normalization$bin_alternative_name <- c("", "1:3", "", "1:2", "", "1:1", "", "2:1", "", "3:1", "")

kable(ratios.normalization,
    format = "html", caption = "Surface areas of the bins", col.names =
        c("Bin", "Surface area", "Bin (left bound)", "Bin (right bound)", "Bin width on x axis", "Bin center", "Integer ratio")
) %>%
    kable_styling("striped", full_width = FALSE)
```

Surface areas of the bins

| Bin | Surface area | Bin (left bound) | Bin (right bound) | Bin width on x axis | Bin center | Integer ratio |
| --- | --- | --- | --- | --- | --- | --- |
| 1 | 0.124255 | 0.0000000 | 0.2352941 | 0.2352941 | 1/4.5 |  |
| 2 | 0.029886 | 0.2352941 | 0.2666667 | 0.0313725 | 1/4 | 1:3 |
| 3 | 0.043699 | 0.2666667 | 0.3076923 | 0.0410256 | 1/3.5 |  |
| 4 | 0.068931 | 0.3076923 | 0.3636364 | 0.0559441 | 1/3 | 1:2 |
| 5 | 0.124617 | 0.3636364 | 0.4444444 | 0.0808081 | 1/2.5 |  |
| 6 | 0.217759 | 0.4444444 | 0.5555556 | 0.1111111 | 1/2 | 1:1 |
| 7 | 0.123828 | 0.5555556 | 0.6363636 | 0.0808081 | 1 - 1/2 |  |
| 8 | 0.068920 | 0.6363636 | 0.6923077 | 0.0559441 | 1 - 1/3 | 2:1 |
| 9 | 0.043541 | 0.6923077 | 0.7333333 | 0.0410256 | 1 - 1/3.5 |  |
| 10 | 0.030120 | 0.7333333 | 0.7647059 | 0.0313725 | 1 - 1/4 | 3:1 |
| 11 | 0.124444 | 0.7647059 | 1.0000000 | 0.2352941 | 1 - 1/4.5 |  |

##### Counting the number of empirical observations in each bin and normalize

Below, we count the number of empirical observations in each bin and
normalize them by dividing by the surface area calculated earlier
multiplied by the total number of observations.

```
# Code off- and on-isochronous
ratios.clean$on_off_isoc <- NA
ratios.clean$on_off_isoc <- factor(ratios.clean$on_off_isoc, levels = c("off", "on", "other"))
ratios.clean$on_off_isoc <- "other"
ratios.clean$on_off_isoc[ratios.clean$bin %in% c(5, 7)] <- "off"
ratios.clean$on_off_isoc[ratios.clean$bin == 6] <- "on"

# Count
ratios.isoc.counts <- dplyr::summarize(dplyr::group_by(ratios.clean, drumming_bout),
    on_isochronous = sum(on_off_isoc == "on"), off_isochronous = sum(on_off_isoc == "off"), other = sum(on_off_isoc == "other"), total_obs = n()
)

# Normalize
ratios.isoc.counts$on_isochronous_norm <- ratios.isoc.counts$on_isochronous / (ratios.isoc.counts$total_obs * ratios.normalization$surface_area[6])
ratios.isoc.counts$off_isochronous_norm <- ratios.isoc.counts$off_isochronous / (ratios.isoc.counts$total_obs *
    (ratios.normalization$surface_area[5] + ratios.normalization$surface_area[7]))
ratios.isoc.counts$other_norm <- ratios.isoc.counts$other / (ratios.isoc.counts$total_obs * (1 - sum(ratios.normalization$surface_area[5:7])))


kable(ratios.isoc.counts,
    format = "html", align = "l",
    caption = "Counts of on- and off-isochronous drumming ratios",
    col.names = c(
        "Drumming bout", "On-isochronous", "Off-isochronous", "Other", "Total observations", "On-isochronous (normalized)",
        "Off-isochronous (normalized)", "Other (normalized)"
    )
) %>%
    kable_styling("striped", full_width = FALSE)
```

Counts of on- and off-isochronous drumming ratios

| Drumming bout | On-isochronous | Off-isochronous | Other | Total observations | On-isochronous (normalized) | Off-isochronous (normalized) | Other (normalized) |
| --- | --- | --- | --- | --- | --- | --- | --- |
| 4 | 11 | 5 | 22 | 38 | 1.3293305 | 0.5296100 | 1.0845854 |
| 5 | 3 | 6 | 5 | 14 | 0.9840499 | 1.7250153 | 0.6690624 |
| 6 | 3 | 5 | 0 | 8 | 1.7220873 | 2.5156473 | 0.0000000 |
| 7 | 7 | 2 | 3 | 12 | 2.6788024 | 0.6708393 | 0.4683437 |
| 8 | 38 | 13 | 2 | 53 | 3.2925442 | 0.9872729 | 0.0706934 |
| 12 | 2 | 1 | 2 | 5 | 1.8368931 | 0.8050071 | 0.7493499 |
| 15 | 13 | 2 | 0 | 15 | 3.9799350 | 0.5366714 | 0.0000000 |
| 16 | 1 | 0 | 1 | 2 | 2.2961163 | 0.0000000 | 0.9366874 |
| 17 | 0 | 1 | 0 | 1 | 0.0000000 | 4.0250357 | 0.0000000 |
| 21 | 9 | 4 | 2 | 15 | 2.7553396 | 1.0733429 | 0.2497833 |
| 22 | 1 | 1 | 0 | 2 | 2.2961163 | 2.0125179 | 0.0000000 |
| 30 | 32 | 16 | 5 | 53 | 2.7726688 | 1.2151051 | 0.1767335 |
| 31 | 14 | 0 | 0 | 14 | 4.5922327 | 0.0000000 | 0.0000000 |
| 34 | 1 | 0 | 1 | 2 | 2.2961163 | 0.0000000 | 0.9366874 |
| 35 | 14 | 0 | 0 | 14 | 4.5922327 | 0.0000000 | 0.0000000 |
| 36 | 12 | 5 | 7 | 24 | 2.2961163 | 0.8385491 | 0.5464010 |
| 37 | 0 | 1 | 0 | 1 | 0.0000000 | 4.0250357 | 0.0000000 |
| 38 | 24 | 10 | 2 | 36 | 3.0614885 | 1.1180655 | 0.1040764 |

The values in the normalized columns represent how much more above
chance level the observed counts are. For example, a value of 1.0 means
that the observed counts are as expected by chance, a value of 2.0 means
that the observed counts are twice what was expected by chance, and so
on.

### Descriptive statistics

#### Before cleaning

##### IOIs descriptives

```
table1(~ percussive_medium + tool_or_limb + percussive_surface + limb + limb_same + display_phase + play_face,
    data = iois, caption = "Descriptive statistics of inter-onset intervals (IOIs) before data cleaning"
)
```

Descriptive statistics of inter-onset intervals (IOIs) before data cleaning

|  | Overall (N=410) |
| --- | --- |
| percussive\_medium |  |
| box | 38 (9.3%) |
| feet | 52 (12.7%) |
| hands | 219 (53.4%) |
| hands+feet | 15 (3.7%) |
| stick1 | 35 (8.5%) |
| stick2 | 45 (11.0%) |
| stick2+feet | 6 (1.5%) |
| tool\_or\_limb |  |
| both | 6 (1.5%) |
| limb | 286 (69.8%) |
| tool | 118 (28.8%) |
| percussive\_surface |  |
| box | 3 (0.7%) |
| box/platform | 1 (0.2%) |
| bushes/ground | 1 (0.2%) |
| bushes/pole | 1 (0.2%) |
| fence | 296 (72.2%) |
| platform | 96 (23.4%) |
| platform/box | 1 (0.2%) |
| platform/bushes | 1 (0.2%) |
| platform/stick2 | 1 (0.2%) |
| pole/bushes | 1 (0.2%) |
| sidefence | 4 (1.0%) |
| stick2 | 3 (0.7%) |
| stick2/platform | 1 (0.2%) |
| limb |  |
| both arms | 1 (0.2%) |
| both arms/left arm | 1 (0.2%) |
| both legs | 6 (1.5%) |
| both legs/left leg | 1 (0.2%) |
| left arm | 315 (76.8%) |
| left arm/both arms | 1 (0.2%) |
| left arm/left leg | 1 (0.2%) |
| left arm/right arm | 2 (0.5%) |
| left arm/right leg | 3 (0.7%) |
| left leg | 7 (1.7%) |
| left leg/left arm | 1 (0.2%) |
| left leg/right arm | 2 (0.5%) |
| left leg/right leg | 15 (3.7%) |
| right arm | 21 (5.1%) |
| right arm/left arm | 2 (0.5%) |
| right arm/left leg | 3 (0.7%) |
| right leg | 10 (2.4%) |
| right leg/both legs | 1 (0.2%) |
| right leg/left arm | 3 (0.7%) |
| right leg/left leg | 14 (3.4%) |
| limb\_same |  |
| False | 50 (12.2%) |
| True | 360 (87.8%) |
| display\_phase |  |
|  | 105 (25.6%) |
| buildup | 6 (1.5%) |
| climax | 229 (55.9%) |
| introduction | 66 (16.1%) |
| letdown | 4 (1.0%) |
| play\_face |  |
| False | 239 (58.3%) |
| False/True | 11 (2.7%) |
| True | 151 (36.8%) |
| True/False | 9 (2.2%) |

##### Onsets descriptives

```
table1(~ percussive_medium + tool_or_limb + percussive_surface + display_phase + play_face,
    data = onsets, caption = "Descriptive statistics of onsets before data cleaning"
)
```

Descriptive statistics of onsets before data cleaning

|  | Overall (N=435) |
| --- | --- |
| percussive\_medium |  |
| box | 39 (9.0%) |
| feet | 56 (12.9%) |
| hands | 228 (52.4%) |
| hands+feet | 16 (3.7%) |
| stick1 | 40 (9.2%) |
| stick2 | 49 (11.3%) |
| stick2+feet | 7 (1.6%) |
| tool\_or\_limb |  |
| both | 7 (1.6%) |
| limb | 300 (69.0%) |
| tool | 128 (29.4%) |
| percussive\_surface |  |
| box | 5 (1.1%) |
| bushes | 2 (0.5%) |
| fence | 308 (70.8%) |
| ground | 1 (0.2%) |
| platform | 109 (25.1%) |
| pole | 1 (0.2%) |
| sidefence | 5 (1.1%) |
| stick2 | 4 (0.9%) |
| display\_phase |  |
|  | 110 (25.3%) |
| buildup | 8 (1.8%) |
| climax | 237 (54.5%) |
| introduction | 75 (17.2%) |
| letdown | 5 (1.1%) |
| play\_face |  |
| False | 271 (62.3%) |
| True | 164 (37.7%) |

##### Ratios descriptives

```
table1(~ percussive_medium + tool_or_limb + percussive_surface + display_phase + play_face,
    data = ratios, caption = "Descriptive statistics of ratios before data cleaning"
)
```

Descriptive statistics of ratios before data cleaning

|  | Overall (N=385) |
| --- | --- |
| percussive\_medium |  |
| box | 37 (9.6%) |
| feet | 48 (12.5%) |
| hands | 210 (54.5%) |
| hands+feet | 14 (3.6%) |
| stick1 | 30 (7.8%) |
| stick2 | 41 (10.6%) |
| stick2+feet | 5 (1.3%) |
| tool\_or\_limb |  |
| both | 5 (1.3%) |
| limb | 272 (70.6%) |
| tool | 108 (28.1%) |
| percussive\_surface |  |
| box | 1 (0.3%) |
| box/box/platform | 1 (0.3%) |
| box/platform/platform | 1 (0.3%) |
| bushes/pole/bushes | 1 (0.3%) |
| fence | 284 (73.8%) |
| platform | 83 (21.6%) |
| platform/box/box | 1 (0.3%) |
| platform/bushes/pole | 1 (0.3%) |
| platform/platform/box | 1 (0.3%) |
| platform/platform/bushes | 1 (0.3%) |
| platform/platform/stick2 | 1 (0.3%) |
| platform/stick2/stick2 | 1 (0.3%) |
| pole/bushes/ground | 1 (0.3%) |
| sidefence | 3 (0.8%) |
| stick2 | 2 (0.5%) |
| stick2/platform/platform | 1 (0.3%) |
| stick2/stick2/platform | 1 (0.3%) |
| display\_phase |  |
| buildup | 5 (1.3%) |
| climax | 221 (57.4%) |
| introduction | 55 (14.3%) |
| introduction/introduction/buildup | 1 (0.3%) |
| letdown | 3 (0.8%) |
| nan/nan/nan | 100 (26.0%) |
| play\_face |  |
| False | 212 (55.1%) |
| False/False/True | 9 (2.3%) |
| False/True/True | 11 (2.9%) |
| True | 138 (35.8%) |
| True/False/False | 4 (1.0%) |
| True/False/True | 2 (0.5%) |
| True/True/False | 9 (2.3%) |

#### After cleaning

##### IOIs descriptives

```
table1(~ percussive_medium + tool_or_limb + percussive_surface + limb + limb_same + display_phase + play_face,
    data = iois.clean,
    caption = "Descriptive statistics of inter-onset intervals (IOIs) after data cleaning"
)
```

Descriptive statistics of inter-onset intervals (IOIs) after data cleaning

|  | Overall (N=313) |
| --- | --- |
| percussive\_medium |  |
| box | 26 (8.3%) |
| feet | 37 (11.8%) |
| hands | 193 (61.7%) |
| stick1 | 22 (7.0%) |
| stick2 | 35 (11.2%) |
| tool\_or\_limb |  |
| limb | 230 (73.5%) |
| tool | 83 (26.5%) |
| percussive\_surface |  |
| fence | 245 (78.3%) |
| platform | 68 (21.7%) |
| limb |  |
| left arm | 276 (88.2%) |
| left leg/right leg | 14 (4.5%) |
| right leg | 10 (3.2%) |
| right leg/left leg | 13 (4.2%) |
| limb\_same |  |
| False | 27 (8.6%) |
| True | 286 (91.4%) |
| display\_phase |  |
|  | 80 (25.6%) |
| climax | 191 (61.0%) |
| introduction | 42 (13.4%) |
| play\_face |  |
| False | 178 (56.9%) |
| True | 135 (43.1%) |

```
iois.descriptives <- ddply(iois.clean, .(), summarize,
    n_obs = length(ioi),
    ioi_min = min(ioi),
    ioi_max = max(ioi),
    ioi_mean = mean(ioi),
    ioi_sd = sd(ioi),
    ioi_median = median(ioi),
    ioi_cov = sd(ioi) / mean(ioi),
    total_ioi_sum = sum(ioi) / 1000
)

kable(iois.descriptives[, 2:ncol(iois.descriptives)],
    format = "html", digits = 2,
    col.names = c("N observations", "Min (ms)", "Max (ms)", "Mean (ms)", "SD (ms)", "Median (ms)", "CV", "Summed duration (s)"),
    caption = "Descriptive statistics of inter-onset intervals (IOIs) after data cleaning"
) %>% kable_styling(bootstrap_options = c("striped", "hover", "condensed", "responsive"))
```

Descriptive statistics of inter-onset intervals (IOIs) after data
cleaning

| N observations | Min (ms) | Max (ms) | Mean (ms) | SD (ms) | Median (ms) | CV | Summed duration (s) |
| --- | --- | --- | --- | --- | --- | --- | --- |
| 313 | 72 | 1365 | 501.27 | 255.35 | 409 | 0.51 | 156.9 |

##### Ratios descriptives

```
table1(~ percussive_medium + tool_or_limb + percussive_surface + display_phase + play_face, data = ratios.clean)
```

|  | Overall (N=309) |
| --- | --- |
| percussive\_medium |  |
| box | 24 (7.8%) |
| feet | 40 (12.9%) |
| hands | 178 (57.6%) |
| hands+feet | 14 (4.5%) |
| stick1 | 23 (7.4%) |
| stick2 | 30 (9.7%) |
| tool\_or\_limb |  |
| limb | 232 (75.1%) |
| tool | 77 (24.9%) |
| percussive\_surface |  |
| fence | 246 (79.6%) |
| platform | 63 (20.4%) |
| display\_phase |  |
| climax | 195 (63.1%) |
| introduction | 39 (12.6%) |
| nan/nan/nan | 75 (24.3%) |
| play\_face |  |
| False | 182 (58.9%) |
| True | 127 (41.1%) |

### Does Toon drum rhythmically?

#### Does Toon drum randomly?

```
ks.test(ratios.clean$ratio, ratios.simulated$ratio)
```

```
    Asymptotic two-sample Kolmogorov-Smirnov test

data:  ratios.clean$ratio and ratios.simulated$ratio
D = 0.21195, p-value = 1.769e-12
alternative hypothesis: two-sided
```

**No, Toon does not drum randomly.**

#### Does Toon drum isochronously?

```
ggplot(ratios.clean, aes(x = ratio, fill = "1")) +
    geom_density(alpha = 0.8, linewidth = 0, show.legend = FALSE) +
    geom_density(aes(x = ratio),
        data = ratios.simulated, color = "goldenrod", linetype = "dashed", linewidth = 2, inherit.aes = FALSE,
        show.legend = FALSE
    ) +
    theme_toon() +
    labs(x = "Ratio", y = "Probability density", title = "Distribution of rhythm ratios") +
    geom_vline(xintercept = breaks, linetype = "dashed", color = "grey") +
    # breaks
    scale_x_continuous(
        breaks = c(0, 0.25, 0.3333, 0.5, 0.6666, 0.75, 1), limits = c(0, 1),
        labels = c("0.0", "0.25\n1:3", "0.33\n1:2", "0.5\n1:1", "0.66\n2:1", "0.75\n3:1", "1.0")
    ) +
    # Make x axis ticks centered
    theme(axis.text.x = element_text(hjust = 0.5))
```

```
ggsave(here("plots", "ratios.pdf"), width = 8, height = 6)
ggsave(here("plots", "ratios.png"), width = 8, height = 6, dpi = 600)
```

The plot suggests absolutely.

Below we compare the proportion of on-isochronous ratios to the
proportion expected by chance (assuming uniformly random IOIs).

We do so by subtracting the surface area of the simulated isochronous
bin from each empirical probability (i.e. \(\frac{\textrm{number of isochronous
observations}}{\textrm{total number of observations}}\)).

Then we test (using a one-sample Wilcoxon signed-rank test) whether
the observed values for each bout are different from 0 (where 0
indicates chance level).

The surface area of the isochronous bin is here calculated using the
formula provided by Jadoul et al. (2024):

\[\frac{1}{2} - \frac{(4b - 5a)^2}{40 (b -
a)^2}\]

```
# Calculate the surface area of the isochronous bin using the formula from Jadoul et al. (2024)
a <- min(iois$ioi)
b <- max(iois$ioi)
surface_area_isochronous <- ((1 / 2) - (((4 * b - 5 * a)^2) / (40 * (b - a)^2)))

# Multiply by two (there's one bin on each side of 0.5 when using this formula)
surface_area_isochronous <- surface_area_isochronous * 2 # 0.2042091

# Subtract surface area of isochronous bin from empirical probabilities
ratios.isoc.counts$isoc_prob <- ratios.isoc.counts$on_isochronous / ratios.isoc.counts$total_obs
ratios.isoc.counts$isoc_prob_normalized <- ratios.isoc.counts$isoc_prob - surface_area_isochronous

write.csv(ratios.isoc.counts, here('data', 'processed', 'isoc_ratio_counts.csv'))
```

```
ggplot(ratios.isoc.counts, aes(y = isoc_prob * 100, x = "", fill = "1")) +
    geom_boxplot(colour = "black", show.legend = FALSE) +
    geom_jitter(width = 0.2, height = 0, show.legend = FALSE) +
    theme_toon() +
    labs(
        y = "Percentage of isochronous drumming ratios per bout", x = "",
        title = "Percentage of isochronous ratios compared to chance level",
        subtitle = "Chance level is based on uniformly randomly distributed IOIs.\nEach dot represents one drumming bout."
    ) +
    geom_hline(yintercept = surface_area_isochronous * 100, linetype = "dashed", linewidth = 2) +
    scale_y_continuous(labels = scales::percent_format(scale = 1))
```

```
# Wilcoxon
wt.isoc <- wilcox.test(ratios.isoc.counts$isoc_prob_normalized, alternative = "two.sided")
wt.isoc
```

```
    Wilcoxon signed rank test with continuity correction

data:  ratios.isoc.counts$isoc_prob_normalized
V = 159, p-value = 0.001454
alternative hypothesis: true location is not equal to 0
```

```
effectsize(wt.isoc)
```

```
r (rank biserial) |       95% CI
--------------------------------
0.86              | [0.64, 0.95]
```

### Does drumming medium change the rhythm?

#### Does the used medium change the duration of the IHIs?

```
iois.clean$tool_or_limb <- as.factor(iois.clean$tool_or_limb)
levels(iois.clean$tool_or_limb) <- c("Limb", "Tool")
ggplot(iois.clean, aes(x = ioi)) +
    geom_histogram(aes(y = ..density.., fill = tool_or_limb), alpha = 0.8, linewidth = 0) +
    geom_density(aes(colour = tool_or_limb), show.legend = FALSE, linewidth = 1.5) +
    theme_toon() +
    labs(x = "Inter-hit interval [ms]", y = "Density", fill = "Percussive implement", title = "Inter-hit intervals by percussive implement")
```

```
ggsave(here("plots", "ioi_duration_by_percussive_medium.pdf"), width = 8, height = 6)
```

This seems to be the case. The drumming is around twice as slow when
using feet, hands, or a box. Below, we compare limb vs. tool.

For the comparison between tool and limb, we combine box and stick,
and feet and hands into ‘tool’ or ‘limb’.

```
ihi.duration.medium <- lm(ioi ~ tool_or_limb, data = iois.clean)
summary(ihi.duration.medium)
```

```
Call:
lm(formula = ioi ~ tool_or_limb, data = iois.clean)

Residuals:
    Min      1Q  Median      3Q     Max 
-566.77 -132.09  -32.09   69.91  933.91 

Coefficients:
                 Estimate Std. Error t value Pr(>|t|)    
(Intercept)        431.09      14.99  28.761   <2e-16 ***
tool_or_limbTool   264.68      29.11   9.093   <2e-16 ***
---
Signif. codes:  0 '***' 0.001 '**' 0.01 '*' 0.05 '.' 0.1 ' ' 1

Residual standard error: 227.3 on 311 degrees of freedom
Multiple R-squared:   0.21, Adjusted R-squared:  0.2075 
F-statistic: 82.69 on 1 and 311 DF,  p-value: < 2.2e-16
```

```
effectsize(ihi.duration.medium)
```

```
# Standardization method: refit

Parameter           | Std. Coef. |         95% CI
-------------------------------------------------
(Intercept)         |      -0.27 | [-0.39, -0.16]
tool or limb [Tool] |       1.04 | [ 0.81,  1.26]
```

**Yes, the duration of the IHI is shorter when using a limb
compared to a tool (an estimated 264.68 ms shorter).**

#### Does the use of limb or tool change drumming variability?

```
var.test(ioi ~ tool_or_limb, data = iois.clean)
```

```
    F test to compare two variances

data:  ioi by tool_or_limb
F = 0.8051, num df = 229, denom df = 82, p-value = 0.2159
alternative hypothesis: true ratio of variances is not equal to 1
95 percent confidence interval:
 0.5538666 1.1348751
sample estimates:
ratio of variances 
         0.8051002
```

**There is no difference in the variability of IOIs between
limb and tool.**

#### Does the use of limb or tool change the amount of isochrony?

```
ratios.clean$tool_or_limb <- as.factor(ratios.clean$tool_or_limb)
levels(ratios.clean$tool_or_limb) <- c("Limb", "Tool")

ggplot(ratios.clean, aes(x = ratio, fill = factor(tool_or_limb, levels = c("Limb", "Tool")))) +
    geom_density(alpha = 0.6, linewidth = 0) +
    geom_density(aes(x = ratio),
        data = ratios.simulated, color = "goldenrod", linetype = "dashed", linewidth = 2, inherit.aes = FALSE,
        show.legend = FALSE
    ) +
    theme_toon() +
    labs(
        x = "Ratio", y = "Probability density", title = "Rhythm ratios by use of tool or limb",
        fill = "Limb vs. tool"
    ) +
    geom_vline(xintercept = breaks, linetype = "dashed", color = "grey") +
    # breaks
    scale_x_continuous(
        breaks = c(0, 0.25, 0.33, 0.5, 0.66, 0.75, 1), limits = c(0, 1),
        labels = c("0.0", "0.25\n1:3", "0.33\n1:2", "0.5\n1:1", "0.66\n2:1", "0.75\n3:1", "1.0")
    )
```

```
ggsave(here("plots", "ratios_by_limb_or_tool.pdf"), width = 8, height = 6)
ggsave(here("plots", "ratios_by_limb_or_tool.png"), width = 8, height = 6, dpi = 600)
```

```
# Remove all observations where the hands_or_feet is not hands or feet
ratios.tool.or.limb <- ratios.clean[ratios.clean$tool_or_limb %in% c("Tool", "Limb"), ]

# Calculate normalized counts
limb.tool.counts <- dplyr::summarize(dplyr::group_by(ratios.tool.or.limb, drumming_bout, tool_or_limb),
    on_isochronous = sum(on_off_isoc == "on"), off_isochronous = sum(on_off_isoc == "off"), other = sum(on_off_isoc == "other"), total_obs = n()
)

# Subtract surface area of isochronous bin from empirical probabilities
limb.tool.counts$isoc_prob <- limb.tool.counts$on_isochronous / limb.tool.counts$total_obs
limb.tool.counts$isoc_prob_normalized <- limb.tool.counts$isoc_prob - surface_area_isochronous

# Wilcoxon
wt.limb.tool <- wilcox.test(
    limb.tool.counts$isoc_prob_normalized[limb.tool.counts$tool_or_limb == "Limb"],
    limb.tool.counts$isoc_prob_normalized[limb.tool.counts$tool_or_limb == "Tool"]
)
wt.limb.tool
```

```
    Wilcoxon rank sum test with continuity correction

data:  limb.tool.counts$isoc_prob_normalized[limb.tool.counts$tool_or_limb == "Limb"] and limb.tool.counts$isoc_prob_normalized[limb.tool.counts$tool_or_limb == "Tool"]
W = 32, p-value = 0.5025
alternative hypothesis: true location shift is not equal to 0
```

```
# plot
ggplot(limb.tool.counts, aes(y = isoc_prob * 100, x = tool_or_limb, fill = tool_or_limb)) +
    geom_boxplot(colour = "black", show.legend = FALSE) +
    geom_jitter(width = 0.2, height = 0, show.legend = FALSE) +
    theme_toon() +
    labs(
        y = "Percentage of isochronous drumming ratios per bout", x = "",
        title = "Percentage of isochronous ratios compared tochance level\nby tool or limb",
        subtitle = "Chance level is based on uniformly randomly distributed IOIs.\nEach dot represents one drumming bout."
    ) +
    geom_hline(yintercept = surface_area_isochronous * 100, linetype = "dashed", linewidth = 2) +
    scale_y_continuous(labels = scales::percent_format(scale = 1)) +
    scale_x_discrete(labels = c("Limb", "Tool"))
```

**There is no statistical difference in the amount of isochrony
when comparing the use of a tool or the limbs.**

### Does the use of hands or feet change the rhythm?

#### Does the use of hands or feet change the duration of the IHI?

```
hands.or.feet.ihi.duration <- lm(ioi ~ hands_or_feet, data = iois.clean)
summary(hands.or.feet.ihi.duration)
```

```
Call:
lm(formula = ioi ~ hands_or_feet, data = iois.clean)

Residuals:
    Min      1Q  Median      3Q     Max 
-592.08 -147.45  -70.45  161.55  885.55 

Coefficients:
                   Estimate Std. Error t value Pr(>|t|)    
(Intercept)          664.08      40.88  16.244  < 2e-16 ***
hands_or_feethands  -184.63      43.53  -4.241 2.94e-05 ***
---
Signif. codes:  0 '***' 0.001 '**' 0.01 '*' 0.05 '.' 0.1 ' ' 1

Residual standard error: 248.7 on 311 degrees of freedom
Multiple R-squared:  0.05467,   Adjusted R-squared:  0.05163 
F-statistic: 17.99 on 1 and 311 DF,  p-value: 2.938e-05
```

```
effectsize(hands.or.feet.ihi.duration)
```

```
# Standardization method: refit

Parameter             | Std. Coef. |         95% CI
---------------------------------------------------
(Intercept)           |       0.64 | [ 0.32,  0.95]
hands or feet [hands] |      -0.72 | [-1.06, -0.39]
```

**Yes, absolutely. Hands are faster than feet, an estimated
184.63 ms faster.**

#### Does the use of hands or feet change drumming variability?

```
print(paste("Standard deviation of IOIs (hands):", sd(iois.clean$ioi[iois.clean$hands_or_feet == "hands"])))
```

```
[1] "Standard deviation of IOIs (hands): 225.335227362217"
```

```
print(paste("Standard deviation of IOIs (feet):", sd(iois.clean$ioi[iois.clean$hands_or_feet == "feet"])))
```

```
[1] "Standard deviation of IOIs (feet): 382.509940203329"
```

```
var.test(ioi ~ hands_or_feet, data = iois.clean)
```

```
    F test to compare two variances

data:  ioi by hands_or_feet
F = 2.8816, num df = 36, denom df = 275, p-value = 1.138e-06
alternative hypothesis: true ratio of variances is not equal to 1
95 percent confidence interval:
 1.837552 4.971204
sample estimates:
ratio of variances 
          2.881557
```

**The IOIs are more variable when using feet compared to
hands.**

#### Does using hands or feet change the amount of isochrony?

```
# Remove all observations where the hands_or_feet is not hands or feet
ratios.hands.or.feet <- ratios.clean[ratios.clean$hands_or_feet %in% c("hands", "feet"), ]

# Calculate normalized counts
hands.feet.counts <- dplyr::summarize(dplyr::group_by(ratios.hands.or.feet, drumming_bout, hands_or_feet),
    on_isochronous = sum(on_off_isoc == "on"), off_isochronous = sum(on_off_isoc == "off"), other = sum(on_off_isoc == "other"), total_obs = n()
)

# Calculate empirical probabilities and subtract surface area of isochronous bin
hands.feet.counts$isoc_prob <- hands.feet.counts$on_isochronous / hands.feet.counts$total_obs
hands.feet.counts$isoc_prob_normalized <- hands.feet.counts$isoc_prob - surface_area_isochronous

# Wilcoxon
wt.hands.feet <- wilcox.test(
    hands.feet.counts$isoc_prob[hands.feet.counts$hands_or_feet == "hands"],
    hands.feet.counts$isoc_prob[hands.feet.counts$hands_or_feet == "feet"]
)
wt.hands.feet
```

```
    Wilcoxon rank sum test with continuity correction

data:  hands.feet.counts$isoc_prob[hands.feet.counts$hands_or_feet == "hands"] and hands.feet.counts$isoc_prob[hands.feet.counts$hands_or_feet == "feet"]
W = 39, p-value = 0.1012
alternative hypothesis: true location shift is not equal to 0
```

```
ggplot(hands.feet.counts, aes(y = isoc_prob * 100, x = hands_or_feet, fill = hands_or_feet)) +
    geom_boxplot(colour = "black", show.legend = FALSE) +
    geom_jitter(width = 0.2, height = 0, show.legend = FALSE) +
    theme_toon() +
    labs(
        y = "Percentage of isochronous drumming ratios per bout", x = "",
        title = "Percentage of isochronous ratios compared to chance level",
        subtitle = "Chance level is based on uniformly randomly distributed IOIs.\nEach dot represents one drumming bout."
    ) +
    geom_hline(yintercept = surface_area_isochronous * 100, linetype = "dashed", linewidth = 2) +
    scale_y_continuous(labels = scales::percent_format(scale = 1))
```

**No, there is no statistical difference in the amount of
isochrony between hands and feet.** **Note that this is
likely due to the small number of observations for feet (in total, 40
ratios for feet, 178 for hands).**

```
ratios.clean$hands_or_feet <- as.factor(ratios.clean$hands_or_feet)
levels(ratios.clean$hands_or_feet) <- c(
    "Feet", "feet/feet/hands", "feet/hands/hands",
    "Hands", "hands/feet/feet", "hands/feet/hands", "hands/hands/feet"
)

ggplot(ratios.clean[ratios.clean$hands_or_feet %in% c("Hands", "Feet"), ], aes(x = ratio, fill = hands_or_feet)) +
    geom_density(alpha = 0.6, linewidth = 0) +
    geom_density(aes(x = ratio),
        data = ratios.simulated, color = "goldenrod", linetype = "dashed", linewidth = 2, inherit.aes = FALSE,
        show.legend = FALSE
    ) +
    theme_toon() +
    labs(
        x = "Ratio", y = "Probability density", title = "Rhythm ratios by hands or feet",
        fill = "Hands vs. feet"
    ) +
    geom_vline(xintercept = breaks, linetype = "dashed", color = "grey") +
    # breaks
    scale_x_continuous(
        breaks = c(0, 0.25, 0.33, 0.5, 0.66, 0.75, 1), limits = c(0, 1),
        labels = c("0.0", "0.25\n1:3", "0.33\n1:2", "0.5\n1:1", "0.66\n2:1", "0.75\n3:1", "1.0")
    )
```

```
ggsave(here("plots", "ratios_by_hands_or_feet.pdf"), width = 8, height = 6)
ggsave(here("plots", "ratios_by_hands_or_feet.png"), width = 8, height = 6, dpi = 600)
```

#### Does the use of alternated limb change the duration of the IHIs?

```
alternation.ihi.duration <- lm(ioi ~ limb_same, data = iois.clean)
summary(alternation.ihi.duration)
```

```
Call:
lm(formula = ioi ~ limb_same, data = iois.clean)

Residuals:
    Min      1Q  Median      3Q     Max 
-479.67 -164.52  -87.52  145.48  868.48 

Coefficients:
              Estimate Std. Error t value Pr(>|t|)    
(Intercept)     551.67      49.13  11.229   <2e-16 ***
limb_sameTrue   -55.15      51.40  -1.073    0.284    
---
Signif. codes:  0 '***' 0.001 '**' 0.01 '*' 0.05 '.' 0.1 ' ' 1

Residual standard error: 255.3 on 311 degrees of freedom
Multiple R-squared:  0.003689,  Adjusted R-squared:  0.0004849 
F-statistic: 1.151 on 1 and 311 DF,  p-value: 0.2841
```

```
effectsize(alternation.ihi.duration)
```

```
# Standardization method: refit

Parameter        | Std. Coef. |        95% CI
---------------------------------------------
(Intercept)      |       0.20 | [-0.18, 0.58]
limb same [True] |      -0.22 | [-0.61, 0.18]
```

**No, there is no difference in the duration of the IHIs
between alternated and non-alternated limb use.**

### Does Toon show limb laterality?

Note that below inlcudes both tool-use and limb-use.

#### Intervals

```
table1(~ limb + limb_same + hands_or_feet, data = iois)
```

|  | Overall (N=410) |
| --- | --- |
| limb |  |
| both arms | 1 (0.2%) |
| both arms/left arm | 1 (0.2%) |
| both legs | 6 (1.5%) |
| both legs/left leg | 1 (0.2%) |
| left arm | 315 (76.8%) |
| left arm/both arms | 1 (0.2%) |
| left arm/left leg | 1 (0.2%) |
| left arm/right arm | 2 (0.5%) |
| left arm/right leg | 3 (0.7%) |
| left leg | 7 (1.7%) |
| left leg/left arm | 1 (0.2%) |
| left leg/right arm | 2 (0.5%) |
| left leg/right leg | 15 (3.7%) |
| right arm | 21 (5.1%) |
| right arm/left arm | 2 (0.5%) |
| right arm/left leg | 3 (0.7%) |
| right leg | 10 (2.4%) |
| right leg/both legs | 1 (0.2%) |
| right leg/left arm | 3 (0.7%) |
| right leg/left leg | 14 (3.4%) |
| limb\_same |  |
| False | 50 (12.2%) |
| True | 360 (87.8%) |
| hands\_or\_feet |  |
| feet | 54 (13.2%) |
| feet/hands | 6 (1.5%) |
| hands | 343 (83.7%) |
| hands/feet | 7 (1.7%) |

**Toon seems to prefer the left hand.**

#### Onsets (limb)

```
table1(~ limb + hands_or_feet, data = onsets)
```

|  | Overall (N=435) |
| --- | --- |
| limb |  |
| both arms | 2 (0.5%) |
| both legs | 8 (1.8%) |
| left arm | 340 (78.2%) |
| left leg | 29 (6.7%) |
| right arm | 28 (6.4%) |
| right leg | 28 (6.4%) |
| hands\_or\_feet |  |
| feet | 65 (14.9%) |
| hands | 370 (85.1%) |

#### Onsets (only left or right)

```
# Count left and right limb onsets based on column 'limb', if the string contains 'left' or 'right'
left.or.right <- onsets %>%
    mutate(left_or_right = ifelse(grepl("left", limb, ignore.case = TRUE), "left", ifelse(grepl("right", limb, ignore.case = TRUE), "right", "both"))) %>%
    group_by(left_or_right) %>%
    dplyr::summarise(n = n())

left.or.right$percentage <- left.or.right$n / sum(left.or.right$n) * 100
kable(left.or.right,
    format = "html", caption = "Counts of left and right limb onsets",
    col.names = c("Limb", "Count", "Percentage of total onsets (%)")
) %>%
    kable_styling("striped", full_width = FALSE)
```

Counts of left and right limb onsets

| Limb | Count | Percentage of total onsets (%) |
| --- | --- | --- |
| both | 10 | 2.298851 |
| left | 369 | 84.827586 |
| right | 56 | 12.873563 |

### Does the used surface change the rhythm?

#### Does the used surface change the duration of the IHIs?

```
ggplot(iois.clean, aes(x = ioi, fill = percussive_surface)) +
    geom_histogram(alpha = 0.5, linewidth = 0) +
    theme_toon() +
    labs(x = "Inter-onset interval [ms]", y = "Probability density", fill = "Percussive surface", title = "IHI by percussive surface")
```

```
ihi.duration.surface <- lm(ioi ~ percussive_surface, data = iois.clean)
summary(ihi.duration.surface)
```

```
Call:
lm(formula = ioi ~ percussive_surface, data = iois.clean)

Residuals:
    Min      1Q  Median      3Q     Max 
-535.56 -156.96  -56.96   92.44  909.04 

Coefficients:
                           Estimate Std. Error t value Pr(>|t|)    
(Intercept)                  455.96      15.38  29.643  < 2e-16 ***
percussive_surfaceplatform   208.60      33.00   6.321 8.99e-10 ***
---
Signif. codes:  0 '***' 0.001 '**' 0.01 '*' 0.05 '.' 0.1 ' ' 1

Residual standard error: 240.8 on 311 degrees of freedom
Multiple R-squared:  0.1139,    Adjusted R-squared:  0.111 
F-statistic: 39.96 on 1 and 311 DF,  p-value: 8.987e-10
```

```
effectsize(ihi.duration.surface)
```

```
# Standardization method: refit

Parameter                     | Std. Coef. |         95% CI
-----------------------------------------------------------
(Intercept)                   |      -0.18 | [-0.30, -0.06]
percussive surface [platform] |       0.82 | [ 0.56,  1.07]
```

**Yes, there is a difference in IHI duration between fence and
platform. The IHI duration is an estimated 208.60 ms slower when
drumming on the platform compared to the fence.**

#### Does the used surface change drumming variability?

```
print(paste("IOI (SD) fence:", sd(iois.clean$ioi[iois.clean$percussive_surface == "fence"])))
```

```
[1] "IOI (SD) fence: 235.041838847254"
```

```
print(paste("IOI (SD) platform:", sd(iois.clean$ioi[iois.clean$percussive_surface == "platform"])))
```

```
[1] "IOI (SD) platform: 260.510084163424"
```

```
var.test(ioi ~ percussive_surface, data = iois.clean)
```

```
    F test to compare two variances

data:  ioi by percussive_surface
F = 0.81403, num df = 244, denom df = 67, p-value = 0.267
alternative hypothesis: true ratio of variances is not equal to 1
95 percent confidence interval:
 0.5423139 1.1709848
sample estimates:
ratio of variances 
         0.8140316
```

**No, there is no difference in the variability of IOIs between
fence and platform.**

#### Does the used surface change the amount of isochrony?

```
# Calculate normalized counts
surface.counts <- dplyr::summarize(dplyr::group_by(ratios.clean, drumming_bout, percussive_surface),
    on_isochronous = sum(on_off_isoc == "on"), off_isochronous = sum(on_off_isoc == "off"), other = sum(on_off_isoc == "other"), total_obs = n()
)

# Calculate empirical probabilities and subtract surface area of isochronous bin
surface.counts$isoc_prob <- surface.counts$on_isochronous / surface.counts$total_obs
surface.counts$isoc_prob_normalized <- surface.counts$isoc_prob - surface_area_isochronous

# Wilcoxon
wt.surface <- wilcox.test(
    surface.counts$isoc_prob[surface.counts$percussive_surface == "fence"],
    surface.counts$isoc_prob[surface.counts$percussive_surface == "platform"]
)
wt.surface
```

```
    Wilcoxon rank sum test with continuity correction

data:  surface.counts$isoc_prob[surface.counts$percussive_surface == "fence"] and surface.counts$isoc_prob[surface.counts$percussive_surface == "platform"]
W = 38.5, p-value = 1
alternative hypothesis: true location shift is not equal to 0
```

**No, there is no difference in the amount of isochrony between
fence and platform.**

### Does the display phase change the rhythm?

#### Does display phase change duration of the IHIs?

```
iois.clean$display_phase <- as.factor(iois.clean$display_phase)
levels(iois.clean$display_phase) <- c("", "Climax", "Introduction")
iois.by.display.phase <- iois.clean[!iois.clean$display_phase == "", ]
ggplot(iois.by.display.phase, aes(x = ioi)) +
    geom_histogram(aes(y = ..density.., fill = display_phase), alpha = 0.8, linewidth = 0) +
    geom_density(aes(colour = display_phase), show.legend = FALSE, linewidth = 1.5) +
    theme_toon() +
    labs(x = "Inter-hit interval [ms]", y = "Density", fill = "Display phase", title = "Inter-hit intervals by display phase")
```

```
ggsave(here("plots", "ihi_duration_by_display_phase.pdf"), width = 8, height = 6)
```

```
print(paste("Mean IHI in introduction:", mean(iois.clean$ioi[iois.clean$display_phase == "Introduction"])))
```

```
[1] "Mean IHI in introduction: 825.238095238095"
```

```
print(paste("Mean IHI in climax:", mean(iois.clean$ioi[iois.clean$display_phase == "Climax"])))
```

```
[1] "Mean IHI in climax: 430.942408376963"
```

```
ihi.duration.display.phase <- lm(ioi ~ display_phase, data = iois.clean[!iois.clean$display_phase == "", ])
summary(ihi.duration.display.phase)
```

```
Call:
lm(formula = ioi ~ display_phase, data = iois.clean[!iois.clean$display_phase == 
    "", ])

Residuals:
    Min      1Q  Median      3Q     Max 
-516.24 -131.94  -51.94   35.06  934.06 

Coefficients:
                          Estimate Std. Error t value Pr(>|t|)    
(Intercept)                 430.94      16.42   26.25   <2e-16 ***
display_phaseIntroduction   394.30      38.67   10.20   <2e-16 ***
---
Signif. codes:  0 '***' 0.001 '**' 0.01 '*' 0.05 '.' 0.1 ' ' 1

Residual standard error: 226.9 on 231 degrees of freedom
Multiple R-squared:  0.3104,    Adjusted R-squared:  0.3074 
F-statistic:   104 on 1 and 231 DF,  p-value: < 2.2e-16
```

```
effectsize(ihi.duration.display.phase)
```

```
# Standardization method: refit

Parameter                    | Std. Coef. |         95% CI
----------------------------------------------------------
(Intercept)                  |      -0.26 | [-0.38, -0.14]
display phase [Introduction] |       1.45 | [ 1.17,  1.73]
```

**Yes, there is a difference in the duration of the IHIs
between display phases. The the duration is an estimated 394.30 ms
shorter in the introduction compared to the climax.**

#### Does display phase change drumming variability?

```
print(paste("IOI (SD) introduction:", sd(iois.clean$ioi[iois.clean$display_phase == "Introduction"])))
```

```
[1] "IOI (SD) introduction: 163.596441240411"
```

```
print(paste("IOI (SD) climax:", sd(iois.clean$ioi[iois.clean$display_phase == "Climax"])))
```

```
[1] "IOI (SD) climax: 238.382469843959"
```

```
var.test(ioi ~ display_phase, data = iois.clean[!iois.clean$display_phase == "", ])
```

```
    F test to compare two variances

data:  ioi by display_phase
F = 2.1232, num df = 190, denom df = 41, p-value = 0.005357
alternative hypothesis: true ratio of variances is not equal to 1
95 percent confidence interval:
 1.261835 3.312727
sample estimates:
ratio of variances 
          2.123249
```

**Yes, drumming in the climax is more variable.**

#### Does display phase change the amount of isochrony?

```
# Calculate normalized counts
display.phase.counts <- dplyr::summarize(dplyr::group_by(ratios.clean, drumming_bout, display_phase),
    on_isochronous = sum(on_off_isoc == "on"), off_isochronous = sum(on_off_isoc == "off"), other = sum(on_off_isoc == "other"), total_obs = n()
)

# Calculate empirical probabilities and subtract surface area of isochronous bin
display.phase.counts$isoc_prob <- display.phase.counts$on_isochronous / display.phase.counts$total_obs
display.phase.counts$isoc_prob_normalized <- display.phase.counts$isoc_prob - surface_area_isochronous

# Wilcoxon
wt.display.phase <- wilcox.test(
    display.phase.counts$isoc_prob[display.phase.counts$display_phase == "introduction"],
    display.phase.counts$isoc_prob[display.phase.counts$display_phase == "climax"]
)
wt.display.phase
```

```
    Wilcoxon rank sum test with continuity correction

data:  display.phase.counts$isoc_prob[display.phase.counts$display_phase == "introduction"] and display.phase.counts$isoc_prob[display.phase.counts$display_phase == "climax"]
W = 26, p-value = 0.8458
alternative hypothesis: true location shift is not equal to 0
```

**No, there is no difference in the amount of isochrony between
introduction and climax.**

```
ratios.clean$display_phase <- as.factor(ratios.clean$display_phase)
levels(ratios.clean$display_phase) <- c("Introduction", "Climax", "nan/nan/nan")


ggplot(ratios.clean[!ratios.clean$display_phase == "nan/nan/nan", ], aes(x = ratio, fill = display_phase)) +
    geom_density(alpha = 0.6, linewidth = 0) +
    geom_density(aes(x = ratio),
        data = ratios.simulated, color = "goldenrod", linetype = "dashed", linewidth = 2, inherit.aes = FALSE,
        show.legend = FALSE
    ) +
    theme_toon() +
    labs(
        x = "Ratio", y = "Probability density", title = "Rhythm ratios by display phase",
        fill = "Display phase"
    ) +
    geom_vline(xintercept = c(0.25, 0.33, 0.5, 0.66, 0.75), linetype = "dashed", color = "grey") +
    # breaks
    scale_x_continuous(
        breaks = c(0, 0.25, 0.33, 0.5, 0.66, 0.75, 1), limits = c(0, 1),
        labels = c("0.0", "0.25\n1:3", "0.33\n1:2", "0.5\n1:1", "0.66\n2:1", "0.75\n3:1", "1.0")
    )
```

```
ggsave(here("plots", "ratios_by_display_phase.pdf"), width = 8, height = 6)
ggsave(here("plots", "ratios_by_display_phase.png"), width = 8, height = 6, dpi = 600)
```

### Does the presence of a play face change the rhythm?

Now, this is a comparison between IOIs where for both hits there was
a play face compared to IOIs where for both hits there was no play
face.

It’s not restricted to bouts that at some point have a play face in
them, but this is in general. I believe it’s difficult to defend why we
would restrict it to bouts that have a play face in them. Also, the
number of observations will be rather low.

```
iois.play.face <- iois.clean[iois.clean$play_face %in% c("True", "False"), ]
iois.play.face$play_face <- as.factor(iois.play.face$play_face)
levels(iois.play.face$play_face) <- c("Play face not present", "Play face present")

ggplot(iois.play.face, aes(x = play_face, y = ioi, fill = play_face)) +
    geom_boxplot(colour = "black", show.legend = FALSE) +
    geom_jitter(width = 0.2, height = 0, show.legend = FALSE) +
    theme_toon() +
    labs(x = "", y = "Inter-hit interval [ms]", fill = "Play face", title = "Inter-hit intervals by play face")
```

```
ggsave(here("plots", "ihi_duration_by_play_face.pdf"), width = 8, height = 6)
```

#### Is play face associated with the duration of the IHIs?

```
iois.play.face <- iois.clean[iois.clean$play_face %in% c("True", "False"), ]
iois.play.face$play_face <- as.factor(iois.play.face$play_face)
levels(iois.play.face$play_face) <- c("Play face not present", "Play face present")

ggplot(iois.play.face, aes(x = ioi)) +
    geom_histogram(aes(y = ..density.., fill = play_face), alpha = 0.8, linewidth = 0) +
    geom_density(aes(colour = play_face), show.legend = FALSE, linewidth = 1.5) +
    theme_toon() +
    labs(x = "Inter-hit interval [ms]", y = "Probability density", fill = "", title = "Inter-hit intervals by play face")
```

```
ggsave(here("plots", "ibi_by_play_face.pdf"), width = 8, height = 6)
```

```
print(paste("Mean IOI with play face", mean(iois.clean$ioi[iois.clean$play_face == "True"])))
```

```
[1] "Mean IOI with play face 373.755555555556"
```

```
print(paste("Mean IOI without play face", mean(iois.clean$ioi[iois.clean$play_face == "False"])))
```

```
[1] "Mean IOI without play face 597.988764044944"
```

```
ihi.duration.play.face <- lm(ioi ~ play_face, data = iois.clean[iois.clean$play_face %in% c("True", "False"), ])
summary(ihi.duration.play.face)
```

```
Call:
lm(formula = ioi ~ play_face, data = iois.clean[iois.clean$play_face %in% 
    c("True", "False"), ])

Residuals:
    Min      1Q  Median      3Q     Max 
-525.99 -162.76   -6.76  131.24  767.01 

Coefficients:
              Estimate Std. Error t value Pr(>|t|)    
(Intercept)     597.99      17.26  34.655  < 2e-16 ***
play_faceTrue  -224.23      26.27  -8.534 6.31e-16 ***
---
Signif. codes:  0 '***' 0.001 '**' 0.01 '*' 0.05 '.' 0.1 ' ' 1

Residual standard error: 230.2 on 311 degrees of freedom
Multiple R-squared:  0.1898,    Adjusted R-squared:  0.1872 
F-statistic: 72.84 on 1 and 311 DF,  p-value: 6.311e-16
```

```
effectsize(ihi.duration.play.face)
```

```
# Standardization method: refit

Parameter        | Std. Coef. |         95% CI
----------------------------------------------
(Intercept)      |       0.38 | [ 0.25,  0.51]
play face [True] |      -0.88 | [-1.08, -0.68]
```

**There definitely seems to be a different duration of the IHIs
when the play face is present. The duration is an estimated 190.19 ms
shorter when the play face is present.**

#### Does the presence of a play face influence drumming variability

```
print(paste("IOI (SD) with play face", sd(iois.clean$ioi[iois.clean$play_face == "True"])))
```

```
[1] "IOI (SD) with play face 90.7600959941957"
```

```
print(paste("IOI (SD) without play face", sd(iois.clean$ioi[iois.clean$play_face == "False"])))
```

```
[1] "IOI (SD) without play face 294.764810753168"
```

```
play.face.var <- var.test(ioi ~ play_face, data = iois.clean[iois.clean$play_face %in% c("True", "False"), ])
play.face.var
```

```
    F test to compare two variances

data:  ioi by play_face
F = 10.548, num df = 177, denom df = 134, p-value < 2.2e-16
alternative hypothesis: true ratio of variances is not equal to 1
95 percent confidence interval:
  7.642764 14.453943
sample estimates:
ratio of variances 
          10.54779
```

```
effectsize(play.face.var)
```

```
F test to compare two variances

Parameter        | Estimate |  df | df (error) |     F |        95% CI |      p
-------------------------------------------------------------------------------
ioi by play_face |    10.55 | 177 |        134 | 10.55 | [7.64, 14.45] | < .001

Alternative hypothesis: true ratio of variances is not equal to 1
```

**When the play face is present the IOIs are less
variable.**

#### Is play face associated with more isochronous rhythms?

```
ratios.play.face <- ratios[ratios$play_face %in% c("True", "False"), ]
ratios.play.face$play_face <- as.factor(ratios.play.face$play_face)
levels(ratios.play.face$play_face) <- c("Play face not present", "Play face present")

ggplot(ratios.play.face, aes(x = ratio)) +
    geom_histogram(aes(y = ..density.., fill = play_face), alpha = 0.8, linewidth = 0) +
    geom_density(aes(colour = play_face), show.legend = FALSE, linewidth = 1.5) +
    theme_toon() +
    labs(x = "Ratio", y = "Density", title = "Rhythm ratios by play face", fill = "Play face") +
    coord_cartesian(xlim = c(0, 1)) +
    theme(legend.title = element_blank())
```

```
ggsave(here("plots", "ratios_by_play_face.pdf"), width = 8, height = 6)
```

```
# Calculate normalized counts
play.face.counts <- dplyr::summarize(dplyr::group_by(ratios.clean, drumming_bout, play_face),
    on_isochronous = sum(on_off_isoc == "on"), off_isochronous = sum(on_off_isoc == "off"), other = sum(on_off_isoc == "other"), total_obs = n()
)

# Calculate empirical probabilities and subtract surface area of isochronous bin
play.face.counts$isoc_prob <- play.face.counts$on_isochronous / play.face.counts$total_obs
play.face.counts$isoc_prob_normalized <- play.face.counts$isoc_prob - surface_area_isochronous

# Wilcoxon
wt.play.face <- wilcox.test(
    play.face.counts$isoc_prob[play.face.counts$play_face == "True"],
    play.face.counts$isoc_prob[play.face.counts$play_face == "False"]
)
wt.play.face
```

```
    Wilcoxon rank sum test with continuity correction

data:  play.face.counts$isoc_prob[play.face.counts$play_face == "True"] and play.face.counts$isoc_prob[play.face.counts$play_face == "False"]
W = 44.5, p-value = 0.5687
alternative hypothesis: true location shift is not equal to 0
```

**No, there is no difference in the amount of isochrony between
play face and no play face.**

### Description of drumming bout 4

#### Descriptives

```
bout.iois <- iois[iois$drumming_bout == 4, ]
bout.ratios <- ratios[ratios$drumming_bout == 4, ]

# IOIs
bout.description <- ddply(bout.iois, .(), summarize,
    n_iois = length(ioi),
    ioi_min_ms = min(ioi),
    ioi_max_ms = max(ioi),
    ioi_mean_ms = mean(ioi),
    ioi_sd_ms = sd(ioi),
    ioi_median_ms = median(ioi),
    ioi_cov = sd(ioi) / mean(ioi),
    total_bout_duration_seconds = sum(ioi) / 1000
)

kable(bout.description[, -1], format = "html", digits = 2, col.names = c(
    "N IOIs", "IOI (min)", "IOI (max)", "IOI (mean)", "IOI (SD)", "IOI (median)",
    "IOI (CV)", "Total duration (s)"
)) %>%
    kable_styling(bootstrap_options = c("striped", "hover", "condensed", "responsive"))
```

| N IOIs | IOI (min) | IOI (max) | IOI (mean) | IOI (SD) | IOI (median) | IOI (CV) | Total duration (s) |
| --- | --- | --- | --- | --- | --- | --- | --- |
| 39 | 141 | 1104 | 676.28 | 356.42 | 918 | 0.53 | 26.38 |

Code 

- Show All Code
- Hide All Code
